# A pore-forming protein drives macropinocytosis to facilitate toad water maintaining

**DOI:** 10.1101/2021.07.31.454564

**Authors:** Zhong Zhao, Zhi-Hong Shi, Chen-Jun Ye, Yun Zhang

**Author notes:** Author to whom correspondence should be addressed: Dr. Yun Zhang, Kunming Institute of Zoology, Chinese Academy of Sciences, 32 East Jiao Chang Road, Kunming 650223, Yunnan, China (PRC); 65198515. These authors contributed equally to this work (Zhong Zhao, Zhi-Hong Shi).

## Abstract

Maintaining water balance is a real challenge for amphibians in terrestrial environments. Our previous studies with toad *Bombina maxima* discovered a secretory aerolysin family pore-forming protein and trefoil factor complex βγ-CAT, which is assembled under tight regulation depending on environmental cues. Here we report an unexpected role for βγ-CAT in toad water maintaining. Deletion of toad skin secretions, in which βγ-CAT is a major component, increased animal mortality under hypertonic stress. βγ-CAT was constitutively expressed in toad osmoregulatory organs, which was inducible under the variation of osmotic conditions. The protein induced and participated in macropinocytosis *in vivo* and *in vitro*. During extracellular hyperosmosis, βγ-CAT stimulated macropinocytosis to facilitate water intake and enhanced exosomes release, which simultaneously regulated aquaporins distribution. Collectively, these findings uncovered that besides membrane integrated aquaporins, a secretory pore-forming protein can facilitate toad water maintaining via macropinocytosis induction and exocytosis modulation, especially in responses to osmotic stress.

## Introduction

Maintaining water balance is a key challenge that amphibians face in the process of transition from water to land (1, 2). In these animals water storage and utilization are enhanced and evaporation is reduced by coordinated functions of the nervous, endocrine and lymphatic systems, and of organs involved in water and salt balance and osmotic regulation, including skin, urinary bladder (UB) and kidney (1, 3). Amphibians are able to absorb water through their skin (1, 2). In addition, water is reabsorbed from the tubular fluid in the kidney and from stored urine in the UB (2, 3). Water channel proteins aquaporins (AQPs) play key roles in transepithelial water absorption/reabsorption in these organs, and in cell volume regulation (4, 5). However, it has rarely been studied in the physiological function of protein components from amphibian skin secretions in regulating of integumental water homeostasis.

Macropinocytosis is a mechanism that mediates bulk uptake and internalization of extracellular fluid and the solutes contained therein, producing endocytic vesicles with diameters of 0.2–5 μm (6, 7). Macropinocytosis is an actin-dependent endocytic pathway mediated by the activation of the Ras and phosphatidylinositol 3-kinase (PI3K)-signaling pathways, which is induced by either endogenous agents such as growth factors, or by invasive microbes (6, 8). This fundamental cellular process has been documented to play various patho-physiological roles in a range of normal and malignant cells, including nutrient acquisition, cell growth, traffic and renewal of membrane components, immune surveillance, entry of pathogens, and cellular motility (6, 9, 10). However, the physiological roles of macropinocytosis, a very ancient form of endocytosis, remain incompletely understood (11, 12).

Pore-forming proteins (PFPs) are usually secretory proteins that exist in a water-soluble monomeric form and oligomerize to form transmembrane pores (channels) (13, 14). Aerolysins are bacterial β-barrel PFPs produced by *Aeromonas* species (13, 14). Interestingly, numerous aerolysin family PFPs (abbreviated af-PFPs, previously referred as Aerolysin-Like Proteins, ALPs) harboring an aerolysin membrane insertion domain fused with other domains have been identified in various animals and plants (15, 16). Our previous studies with the skin secretions of the toad *Bombina maxima* discovered an interaction network among af-PFPs and trefoil factors (TFFs) (17). BmALP1, an af-PFP from the toad, can be reversibly regulated between the active and inactive forms, in which its paralog BmALP3 is a negative regulator depending on environmental oxygen tension (18). Specifically, BmALP1 interacts with BmTFF3 to form a membrane active PFP complex named βγ-CAT (19–21), in which BmTFF3 acts as an extracellular chaperon that stabilizes the BmALP1 monomer and delivers this PFP to its proper membrane targets (18, 21, 22).

This secretory PFP complex βγ-CAT targets gangliosides and sulfatides in cell membranes in a double-receptor binding model (22). Then the BmALP1 subunit is endocytosed and this PFP oligomerizes to form channels on endolysosomes, modulating the contents and biochemical properties of these intracellular organelles (22–25). Thus, βγ-CAT is a secretory endolysosome channel (SELC) protein, representing a hitherto unknown SELC pathway (17). Depending on cell contexts and surroundings, the cellular effects of SELC protein βγ-CAT have been proposed to facilitate the toad in the sense and uptake of environmental materials (like nutrients and antigens) and vesicular transport, while maintaining mucosal barrier function and fulfilling immune defense (17). Accordingly, the roles of βγ-CAT in immune defense have been first documented (21, 23–26).

Amphibian skin is a major organ responsible for water acquisition and maintaining (1, 3), and βγ-CAT is a major component of *B. maxima* skin secretions (18). In the present study, we found that the expression and localization of βγ-CAT in toad *B. maxima* are related to environmental osmotic conditions, and that this protein is able to counteract cellular dehydration under extracellular hyperosmosis. βγ-CAT stimulated and participated in cell macropinocytosis and exosome release, promoting water and Na^+^ uptake and regulating AQP localization. Collectively, these results revealed that a secretory PFP can drive water acquisition and maintaining.

## Results

### βγ-CAT is involved in responses to osmotic stress

Amphibian skin has played a principal role in water balance during evolutionary adaptation to land environments (1, 2). Accordingly, when exposed to hypertonic Ringer’s solution, toads (*B. maxima*) rapidly lost weight, which was gradually recovered when the animal was then placed into isotonic Ringer’s solution (Fig. 1*A*). This suggests that the toad transports water through its skin under conditions of osmotic stress. Skin secretions play pivotal roles in the physiological functions of amphibian skin (2). Furthermore, βγ-CAT is a major proteinaceous component of skin secretions in toad *B. maxima* (18). Two subunits of βγ-CAT together were estimated to account for about 50% of proteins in *B. maxima* skin secretions (Fig. S1*A*). We speculated that toad skin secretions, especially βγ-CAT, might play an active role in water balance. Osmotic stress experiments were conducted to test the possible functions of βγ-CAT-containing skin secretions in maintenance of water balance in the toad. Toads *B. maxima* lost 20% of their body weight and 50% of the animals died when they were placed in the hypertonic Ringer’s solution for 24 hours (Fig. 1*B* and Fig. S1*B*). Toad skin secretions were depleted by electro-stimulation 30 minutes before the animals were placed in different osmotic solutions. Although no toad deaths were recorded after electro-stimulation in the isotonic Ringer’s solution, significant mortality occurred in the hypertonic Ringer’s solution (Fig. 1*B*). Therefore, it appears that skin secretions are critical for the toad to cope with hypertonic environments. To further investigate the possible involvement of βγ-CAT in the toad’s water balance, real-time fluorescent quantitative PCR was used to detect the expression of two βγ-CAT subunits in the skin, UB and kidney of toad *B. maxima* during dehydration and weight recovery (water absorption after dehydration). It was found that βγ-CAT α-subunit expression was upregulated in the skin and kidney during weight recovery via water uptake after dehydration (Fig. 1*C* and Fig. S1*C*). In contrast, the expression of βγ-CAT β-subunit mRNA was upregulated in the toad skin during dehydration or water absorption (Fig. 1*D*), but not in the kidney (Fig. S1*D*). Interestingly, in the UB, the mRNA levels of the both subunits were increased when toads were exposed to hypertonic Ringer’s solution, and returned to the normal level when toads were subsequently placed in isotonic Ringer’s solution (Fig. 1 *E* and *F*). Therefore, βγ-CAT expression is associated with changes to external osmotic conditions.

**Fig. 1.**
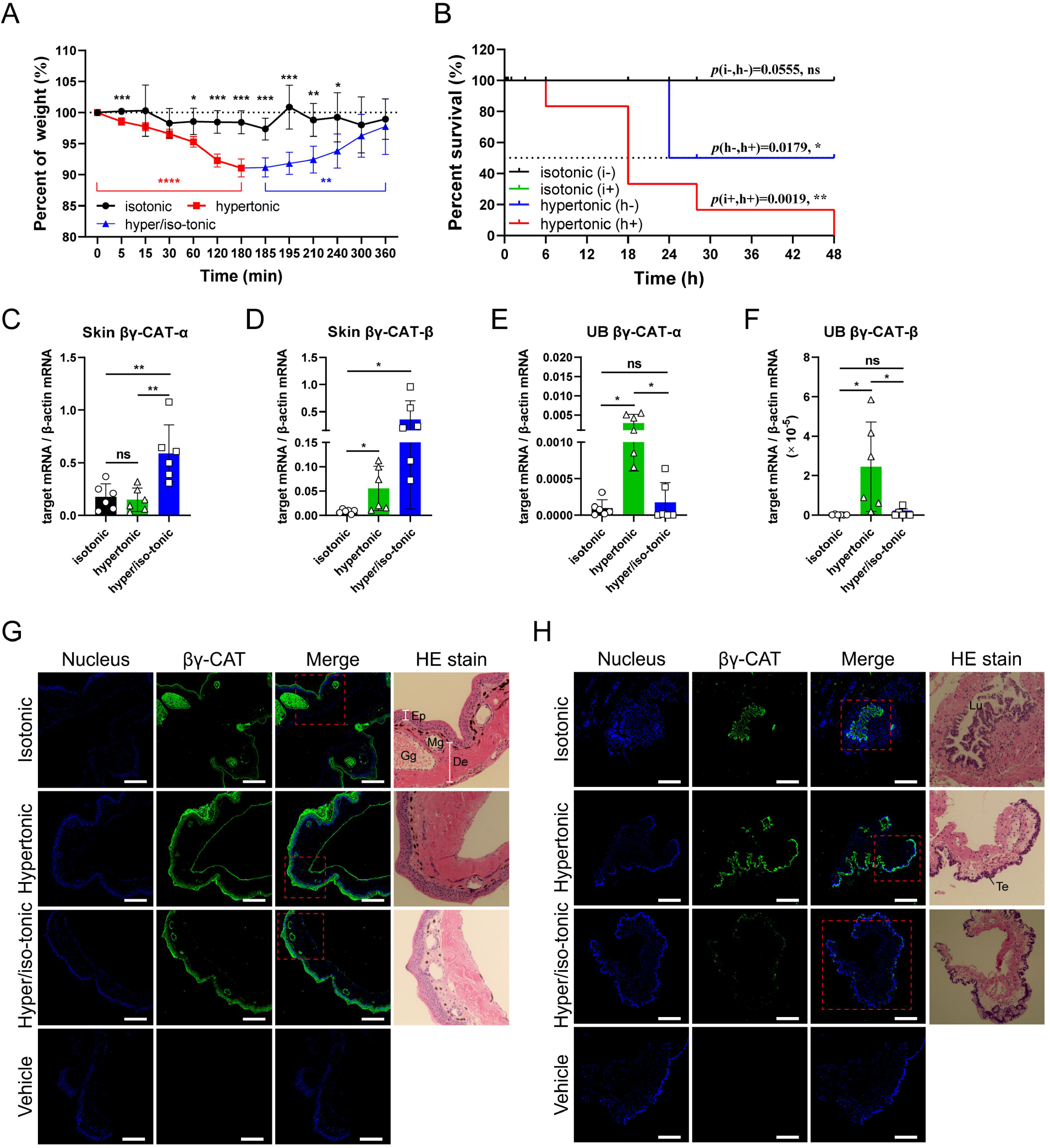
βγ-CAT is involved in responses to osmotic stress. (*A*) Toad weight changes were measured after placing them in isotonic, hypertonic and hypertonic/isotonic Ringer’s solutions. Initial weight of toads is 19 ± 5 g (*n* = 5). (*B*) Survival rates of toads in isotonic (i) and hypertonic (h) Ringer’s solution were determined after 48 hours. The animals were placed in each of the solutions after 30 min with (+) or without (-) electro-stimulation to delete toad skin secretions (*n* = 6). (*C*-*F*) The expression of βγ-CAT subunits in toad skin and UB were analyzed by real-time fluorescent quantitative PCR (*n* = 6). The animals were placed in isotonic or hypertonic Ringer’s solution for 3 hours before the samples were collected. In the hypertonic/isotonic group, the toad was first placed in the hypertonic solution for 3 hours, moved to isotonic solution for a further 3 hours, and then the sample was collected. (*G, H*) Following placement of toads in isotonic, hypertonic or hypertonic/isotonic Ringer’s solution as described above, the localization and expression of βγ-CAT in the toad skin (*G*) and UB (*H*) tissues were analyzed by immunohistofluorescence (IHF). Ep, epidermis; De, dermis; Mg, mucous gland; Gg, granular gland; Lu, lumen; Te, transitional epithelium, Scale bars, 500 μm. All data represent the mean ± SD of two independent replicated experiments. \**P* < 0.05 and \*\**P* < 0.01 by the Gehan-Breslow-Wilcoxon test in survival rate analysis. ns (*P* ≥ 0.05); \**P* < 0.05, \*\**P* < 0.01 and \*\*\**P* < 0.001 by unpaired *t* test in other experiments. All data are representative of at least two independent experiments in *G, H*. See also Fig. S1.

In addition, we analyzed the expression and localization of βγ-CAT protein under the same treatment conditions via immunohistofluorescence (IHF). βγ-CAT was mainly concentrated within the epidermis and glands of the skin, and it was obviously upregulated in the epidermis and basement membrane during osmotic stress (Fig. 1*G*). βγ-CAT was predominantly localized within the transitional epithelium in the UB. The expression of βγ-CAT protein in toad UB was upregulated in hypertonic Ringer’s solution and downregulated when the toad was then placed into isotonic Ringer’s solution (Fig. 1*H*). Collectively, these results revealed an involvement of βγ-CAT in the toad’s responses to osmotic stress.

### βγ-CAT counteracts cell dehydration under extracellular hyperosmosis

βγ-CAT is a vital functional protein of *B. maxima*, and its constitutive expression can be detected in various toad tissues (18, 23), including skin secretions and epithelial cells from the skin and UB, as well as kidney and peritoneal cells. Endogenous secretion of βγ-CAT in these toad-derived cells was further analyzed by a hemolysis assay, a sensitive method to detect the presence of biologically active βγ-CAT (18, 19). All the media of these cultured toad cells showed potent hemolytic activity, which was totally inhibited by anti-βγ-CAT antibodies, confirming the presence of secreted βγ-CAT (Fig. S2*A*). In addition, we measured the cytotoxicity of βγ-CAT to *B. maxima* cells. Previously, it was reported that βγ-CAT showed no cytotoxicity to toad peritoneal cells at dosages up to 400 nM (23). The present study further determined that cytotoxicity of βγ-CAT to skin and UB epithelial cells and kidney cells of *B. maxima* occurred only when its concentration reached 2 μM (Fig. S2*B*), a level much higher than physiological concentrations (20–100 nM, as determined in the toad peritoneum) (23). In contrast, mammalian cells are much more sensitive to βγ-CAT. Although 10 nM of βγ-CAT showed no cytotoxicity to MDCK, Caco-2 and T24 cells, the protein caused significant cell death when dosages used were higher than 50–100 nM (Fig. S2*C*). Thus, in all subsequent experiments concerning the addition of purified βγ-CAT to cells, the protein dosages used were: epithelial cells from toad skin, 100 nM; UB epithelial cells, 50 nM; peritoneal cells, 50 nM; mammalian MDCK, 10 nM; Caco-2, 10 nM; and T24, 5 nM.

To further explore the potential role of βγ-CAT in water transport, we investigated whether the protein is able to counteract cell dehydration under extracellular hyperosmosis. We confirmed the action of βγ-CAT on toad UB epithelial cells, mammalian MDCK, Caco-2 and T24 cells, as determined by the formation of βγ-CAT oligomers after treatment with the protein (Fig. 2*A*). βγ-CAT oligomers were detected in toad UB epithelial cells without the addition of the protein, possibly caused by endogenously secreted βγ-CAT (Fig. 2*A*). The addition of purified βγ-CAT further increased the concentration of oligomers of the protein in the UB epithelial cells (Fig. 2*A*). Then, we analyzed the electrophysiological characteristics of βγ-CAT oligomers on outside-out patches of HEK293 cells. Currents were elicited by 500 ms ramp protocol between -100 mV to +100 mV every 2 s from a holding potential of 0 mV. βγ-CAT induced macroscopic currents were recorded and normalized, which has the characteristic of inward rectification (Fig. 2*B*). These proved that βγ-CAT could form transmembrane pores after oligomerization on the membrane and mediate ion flow. We further investigated ion selectivity of βγ-CAT channels through ion replacement experiments. Asymmetric 150:15 mM NaCl solutions left-shifted the reversal potential from 0 mV to -61.2 mV, which is very close to the theoretical equilibrium potential of Na^+^, indicating that the βγ-CAT channels are permeable to Na^+^ but not to Cl^-^. (Fig. 2*C*). These results demonstrated that the βγ-CAT channels allowed Na^+^ flow.

**Fig. 2.**
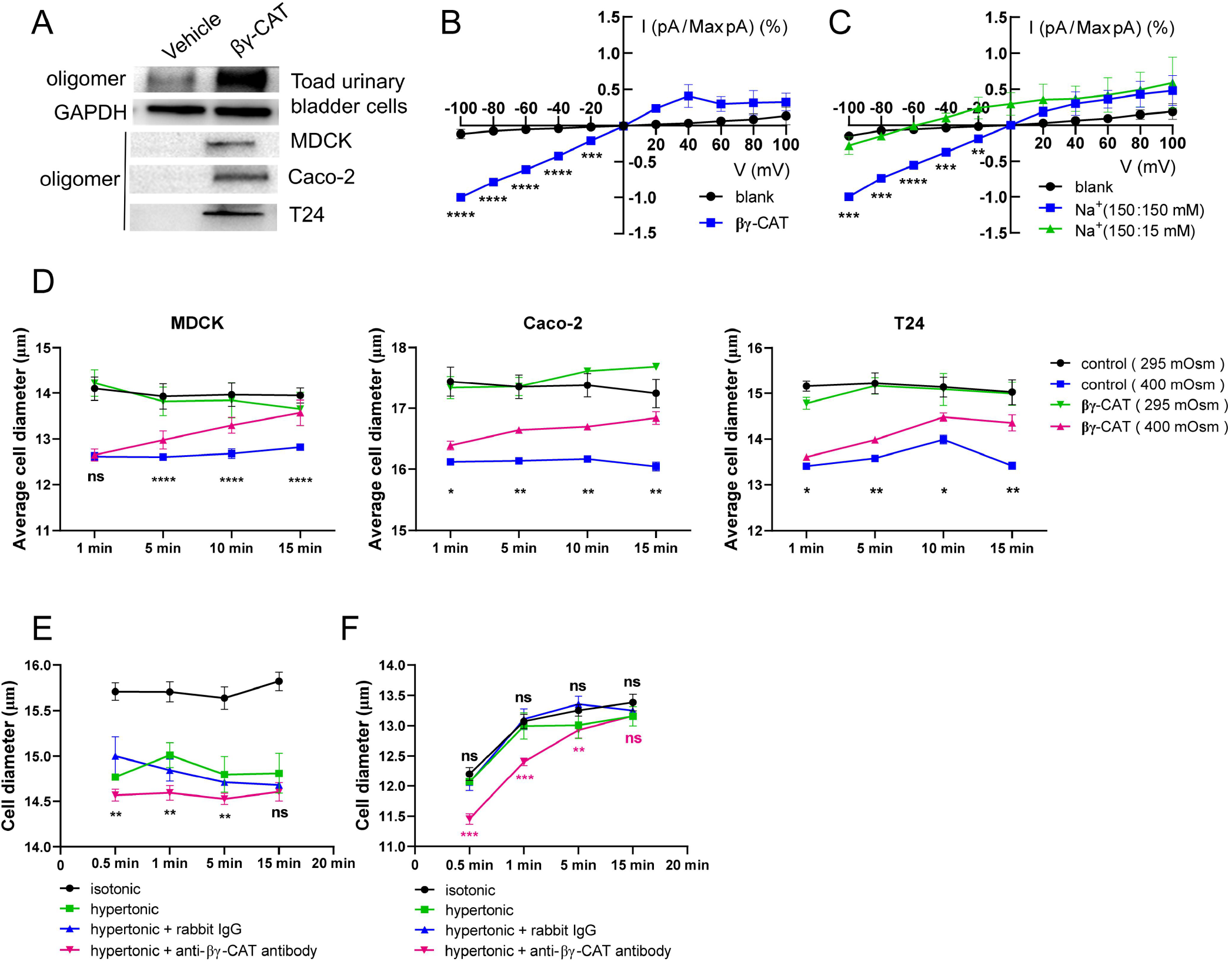
βγ-CAT counteracts cellular dehydration under extracellular hyperosmosis. (*A*) Various mammalian and toad cells were first treated with or without βγ-CAT for 15 minutes. The appearance of βγ-CAT oligomers in the treated cells was determined by western blotting. (*B*) Normalized current-voltage curves of channels formed by 100 nM βγ-CAT on HEK293 cells. Currents were elicited by 500 ms ramp protocol between -100 mV to +100 mV every 2 s from a holding potential of 0 mV (*n* = 3). (*C*) Normalized current-voltage curves of βγ-CAT channels in indicated solutions (pipette/bath) (*n* = 3). Na^+^(150 : 15 mM) represents the 150 mM Na^+^ in the pipette and 15 mM Na^+^ in the bath. (*D*) Diameter changes of MDCK, Caco-2 and T24 cells in isotonic (295 mOsm) or hypertonic (400 mOsm) PBS in the presence or absence of βγ-CAT as determined by using a cell counter. Digested cells were first suspended in PBS for 15 minutes before they were used in this experiment. (*E, F*) Diameter changes of toad UB epithelial cells in isotonic (black) and hypertonic (green) Ringer’s solution in the presence of 50 μg/mL rabbit antibody (blue) or anti-βγ-CAT antibody (magenta). In (*F*), the cells were first treated with 0.3 mM HgCl_2_ for 10 minutes before cell diameter measurement with a cell counter. In all experiments shown in Figure 2, βγ-CAT dosages used were 10 nM for MDCK and Caco-2, 5 nM for T24 and 50 nM for toad UB epithelial cells. The bars represent the mean ± SD of triplicate samples in *B*-*E*. The bars represent the mean ± SD of three independent replicates in *F*. ns (*P* ≥ 0.05), \**P* < 0.05, \*\**P* < 0.01, \*\*\**P* < 0.001 and \*\*\*\**P* < 0.0001 by the unpaired *t* test. All data are representative of at least two independent experiments. See also Fig. S2.

On this basis, we explored the potential role of βγ-CAT when cells were challenged by extracellular hyperosmosis using hypertonic solution. We observed that βγ-CAT promoted volume recovery of MDCK, Caco-2 and T24 cells in hypertonic solution (Fig. 2*D*), revealing the capacity of the protein to facilitate water uptake under hyperosmotic conditions. In consistency with those observations, immune-depletion of endogenous βγ-CAT further decreased the cell diameters of toad UB epithelial cells in hypertonic Ringer’s solution relative to those in the absence of anti-βγ-CAT antibodies (Fig. 2*E*). It is well documented that AQPs can mediate the rapid cellular flow of water (27). Thus, it is necessary to clarify whether the cell volume recovery by water uptake mediated by βγ-CAT under osmotic stress was associated with AQP function. Sequence alignment and evolutionary analysis of *B. maxima* AQPs based on a toad UB transcriptome revealed that AQP1, AQP2, AQP3, AQP5 and AQP11 were present in *B. maxima* UB tissue (Fig. S2*D*). Because the mercury-sensitive sites of *B. maxima* AQPs (BmAQPs) were evolutionarily conserved (Fig. S2*E*), HgCl_2_ was used to inhibit BmAQPs function. Cell volume was unchanged in hypertonic Ringer’s solution after blocking BmAQPs with HgCl_2_. Conversely, cell volume recovery of toad UB epithelial cells via water uptake was clearly reduced after immunodepletion of endogenous βγ-CAT (Fig. 2*F*). Taken together, these results revealed the capacity of βγ-CAT to promote cell volume recovery by stimulating water acquisition during extracellular hyperosmosis, and indicated that the effect was independent of water flow via plasma membrane AQPs.

### βγ-CAT promotes macropinocytosis

βγ-CAT participates in cell volume regulation, suggesting its involvement in water transport. We next analyzed the possible cellular mechanism behind this interesting phenomenon. Water acquisition can be rapidly realized through AQPs and/or macropinocytosis (12, 27, 28). Previously, in a murine dendritic cell (DC) model, preliminary data suggested that βγ-CAT might enhance pinocytosis (25). In the present study, we carefully studied the ability of this protein to stimulate and participate in macropinocytosis in various types of *B. maxima* cells. First, immunoelectron microscopy (IEM) revealed that βγ-CAT was localized in cellular pseudopodia and macropinosomes with diameters up to 300 nm formed by macropinocytosis in toad skin and UB tissues (Fig. 3*A*). βγ-CAT was also present in the intercellular spaces of epithelial cells (Fig. 3*A*). Lucifer Yellow (LY) and 70-kDa dextran are fluid-phase specific markers for macropinocytosis (29, 30). In epithelial cells obtained from toad skin and UB, and in toad kidney or peritoneal cells, immunodepletion of endogenous βγ-CAT by anti-βγ-CAT antibodies greatly decreased internalization of LY and FITC-dextran, relative to cells observed in the presence of control rabbit IgG (Fig. 3*B*, 3*C*, S3*A* and S3*B*). Furthermore, addition of βγ-CAT to cells from toad skin and UB, and mammalian MDCK and T24 cells substantially enhanced the internalization of LY and FITC-dextran (Fig. 3*D*, 3*E*, S3*C* and S3*D*). Confocal microscopy showed that βγ-CAT was endocytosed and located in the macropinosomes of toad UB epithelial cells (Fig. S3*E*), peritoneal cells (Fig. S3*F*), and mammalian MDCK cells (Fig. S3*G*). In addition, we found that total Na^+^ concentrations in toad UB epithelial cells and MDCK treatment with βγ-CAT were 3.5 and 1.1 times of that of the control, respectively (Fig. 3*F*).

**Fig. 3.**
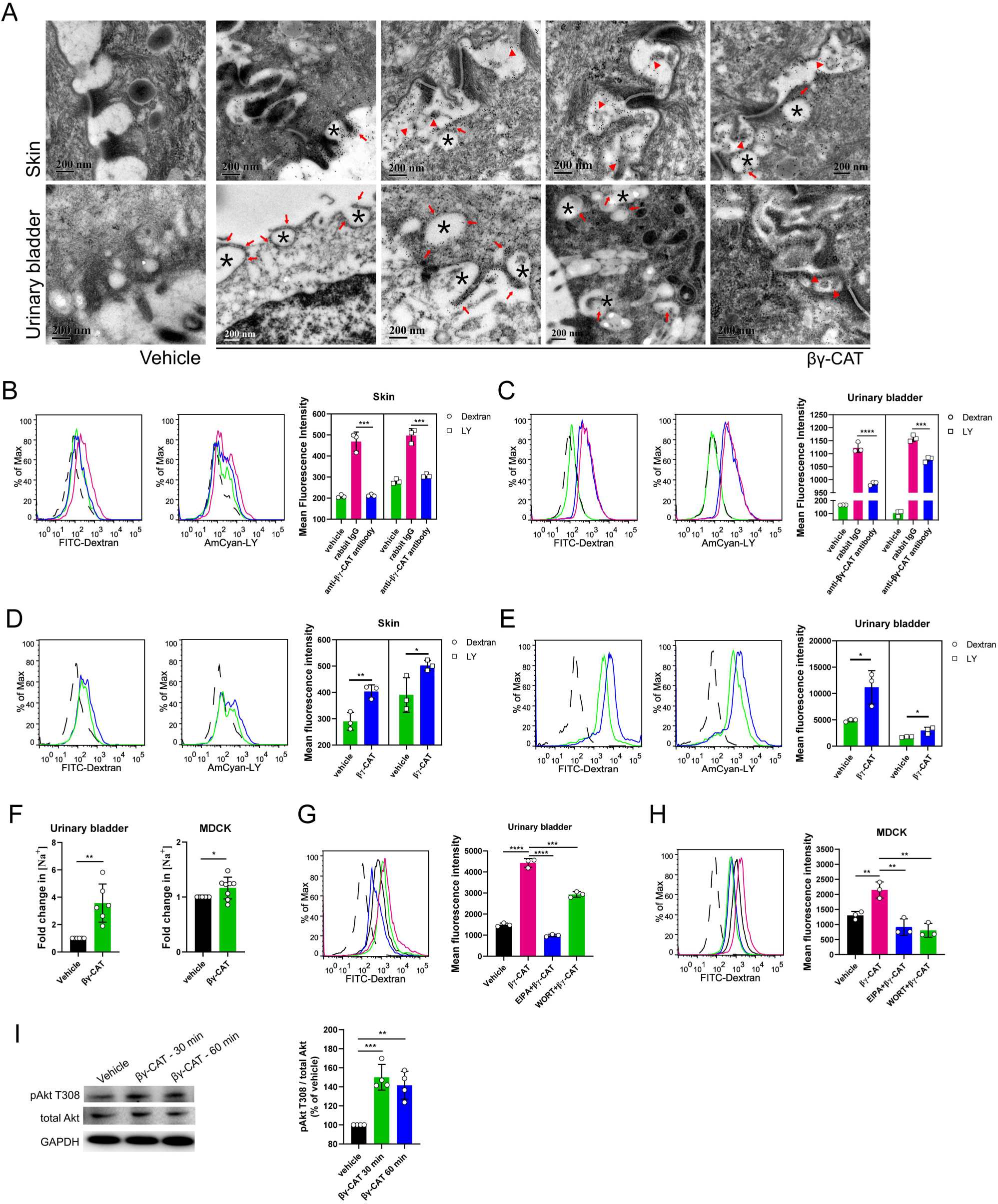
βγ-CAT promotes macropinocytosis. (*A*) Ultrastructural localization of βγ-CAT in toad skin and UB tissues as analyzed by IEM. Vehicle (rabbit IgG control). Endocytic vesicles formed by macropinocytosis (black asterisks), and distribution of βγ-CAT on vesicles (red arrows) and intercellular spaces (red triangles). (*B, C*) Immunedepletion of endogenous βγ-CAT decreased macropinocytosis. Toad skin (*B*) and UB (*C*) epithelial cells were incubated with 50 μg/mL anti-βγ-CAT antibody to immunodeplete endogenous βγ-CAT for 30 minutes. The mean fluorescence intensity was determined by flow cytometry with 100 μg/mL of 70 kDa FITC-label dextran and Lucifer Yellow (LY) for 30 minutes. Rabbit IgG (antibody control). Vehicle (antibody absent control). (*D, E*) The addition of purified βγ-CAT augmented macropinocytosis. The mean fluorescence intensity of LY and FITC-label dextran in toad skin (d) and UB (e) epithelial cells was determined by flow cytometry with or without additional 100 nM or 50 nM βγ-CAT, respectively. (*F*) Fold changes of [Na^+^] in toad UB epithelial cells (left, *n* = 6) and MDCK (right, *n* = 8) with and without the addition of 50 nM or 10 nM βγ-CAT for 3 hours, respectively. (*G, H*) The effect of inhibitors on macropinocytosis induced by βγ-CAT. Toad UB epithelial cells (*G*) and MDCK cells (*H*) were incubated with and without 100 μM EIPA or 20 μM WORT for 1 hour. Then the cells were cultured with 100 μg/mL FITC-label dextran with 50 nM (toad UB cells) or 10 nM (MDCK cells) βγ-CAT for 30 minutes. (*I*) Akt phosphorylation in response to 10 nM βγ-CAT in MDCK cells for 30 or 60 minutes as determined by western blotting (left), and bands were semiquantified with ImageJ (right). The black dotted line refers to the blank control, and data represent the mean ± SD of triplicate samples in *B*-*E, G, H*. Data represent the mean ± SD of at least three independent replicates in *F* and *I*. \**P* < 0.05, \*\**P* < 0.01, \*\*\**P* < 0.001 and \*\*\*\**P* < 0.0001 by the unpaired *t* test. All data are representative of at least two independent experiments. See also Fig. S3.

Next, the molecular mechanism of macropinocytosis induced by βγ-CAT was investigated using pharmacological agents. 5-(N-ethyl-N-isopropyl) amiloride (EIPA, a Na^+^/H^+^-exchanger inhibitor) and wortmannin (WORT, a PI3K inhibitor) are commonly used to inhibit macropinocytosis (31, 32). We found that EIPA and WORT reduced macropinocytosis induced by βγ-CAT both in toad UB epithelial cells and mammalian MDCK cells (Fig. 3 *G* and *H*). Moreover, βγ-CAT stimulated an increase in Akt phosphorylation in MDCK cells (Fig. 3*I*). Taken together, these results demonstrated that βγ-CAT, an endogenously secreted complex of a PFP and a TFF, stimulates and participates in macropinocytosis in diverse cells of *B. maxima* as well as in various mammalian cells.

### βγ-CAT in AQP regulation

AQPs and ion flux are involved in water transport and volume regulation of toad UB cells under stimulation by multiple hormones (33–35). Internalization by macropinocytosis is unselective, whereas transitional epithelial cells are polar. Therefore, we assessed the relationship between βγ-CAT-stimulated macropinocytosis and BmAQP2. We observed that the location of βγ-CAT and BmAQP2 in toad UB epithelial cells differed between isotonic, hypertonic and ‘hypertonic/isotonic’ (hypertonic followed by return to isotonic) Ringer’s solutions (Fig. 4*A*). When toads were exposed to isotonic Ringer’s solution, βγ-CAT and BmAQP2 were located together and were distributed on both the apical and basal sides of the transitional epithelium in toad UB. In hypertonic Ringer’s solution, βγ-CAT and BmAQP2 shifted their location to the basal or lateral sides of the UB transitional epithelium. In contrast, when the toads were returned to isotonic Ringer’s solution from hypertonic Ringer’s solution, βγ-CAT and BmAQP2 began to migrate to the apical side of the transitional epithelium. This observation suggests that macropinocytosis induced by βγ-CAT is involved in the internalization and transport of BmAQP2 in the UB. In MDCK, intracellular colocalization of βγ-CAT and AQP2 or AQP3 was also observed (Fig. 4 *B* and *C*). These results indicate that macropinocytosis stimulated by βγ-CAT plays a role in the regulation of AQPs under various osmotic conditions, revealing another aspect of this protein in the modulation of toad water balance.

**Fig. 4.**
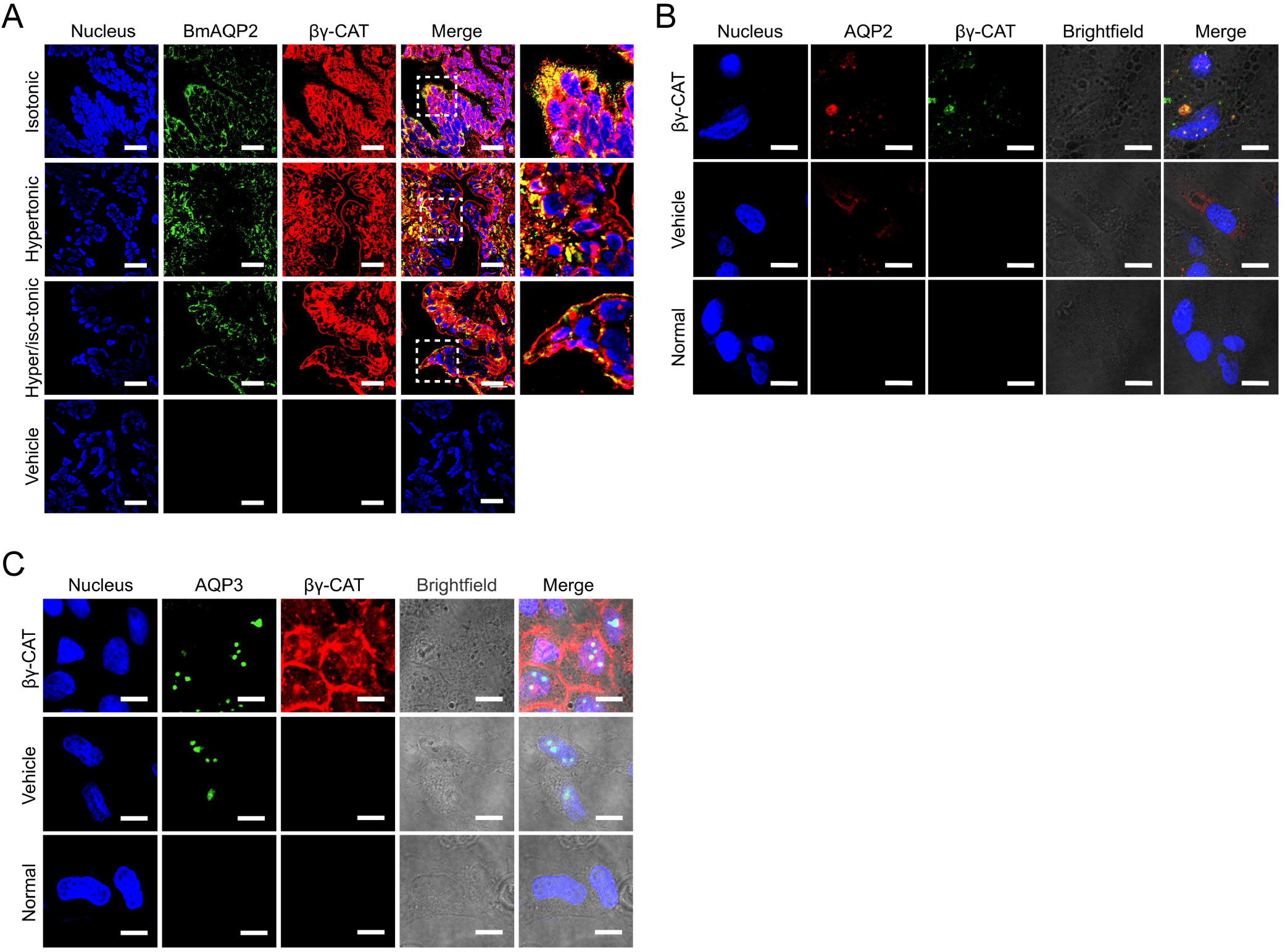
βγ-CAT in AQP regulation. (*A*) Colocalization of βγ-CAT and BmAQP2 in the UB tissue of *B. maxima* after the animals were placed in isotonic, hypertonic or hypertonic/isotonic Ringer’s solution for 3 hours as analyzed by immunohistofluorescence (Scale bars, 25 μm). (*B, C*) Intracellular colocalization of βγ-CAT and AQP2 (*B*) or AQP3 (*C*) in MDCK cells with or without the treatment of 10 nM βγ-CAT for 15 minutes as determined by immunofluorescence (Scale bars, 10 μm). All data are representative of at least two independent experiments.

### βγ-CAT enhances exosome release

The secretion of exosomes is an important aspect of cell exocytosis that can be viewed as a means of selective transport of materials among cells and a mode of intercellular communications (36). Because βγ-CAT promotes macropinocytosis, we further explored the cellular fate and role of βγ-CAT-containing vesicles. βγ-CAT was readily detected by IEM in both the multivesicular bodies (MVBs) and the intraluminal vesicles (ILVs) of *B. maxima* UB epithelial cells (Fig. 5*A*). Furthermore, it was noted that the survival rate of toad UB epithelial cells was greater than 98% after *in vitro* culture for 3 hours at room temperature (Fig. S4*A*). Thus, exosomes secreted by these cells were collected and their shapes were observed by TEM (Fig. 5*B*). The exosome markers CD63, TSG101 and flotillion-1 were detected in the collected exosomes and were substantially augmented in UB epithelial cells treated by the addition of 50 nM purified βγ-CAT (Fig. 5*C*). Nanoparticle tracking analysis (NTA) revealed that immunodepletion of endogenous βγ-CAT greatly attenuated exosome release from toad UB epithelial cells and peritoneal cells (Fig. 5*D* and Fig. 4*B*), while the addition of βγ-CAT substantially augmented exosome release from these cells (Fig. 5*E* and Fig. S4*C*). It is noteworthy that none of these treatments changed the average exosome diameter. Similar results were also obtained in mammalian MDCK and T24 cells (Fig S4 *D* and *E*). Collectively, these results showed that βγ-CAT promotes the production and release of exosomes in both toad and mammalian cells.

**Fig. 5.**
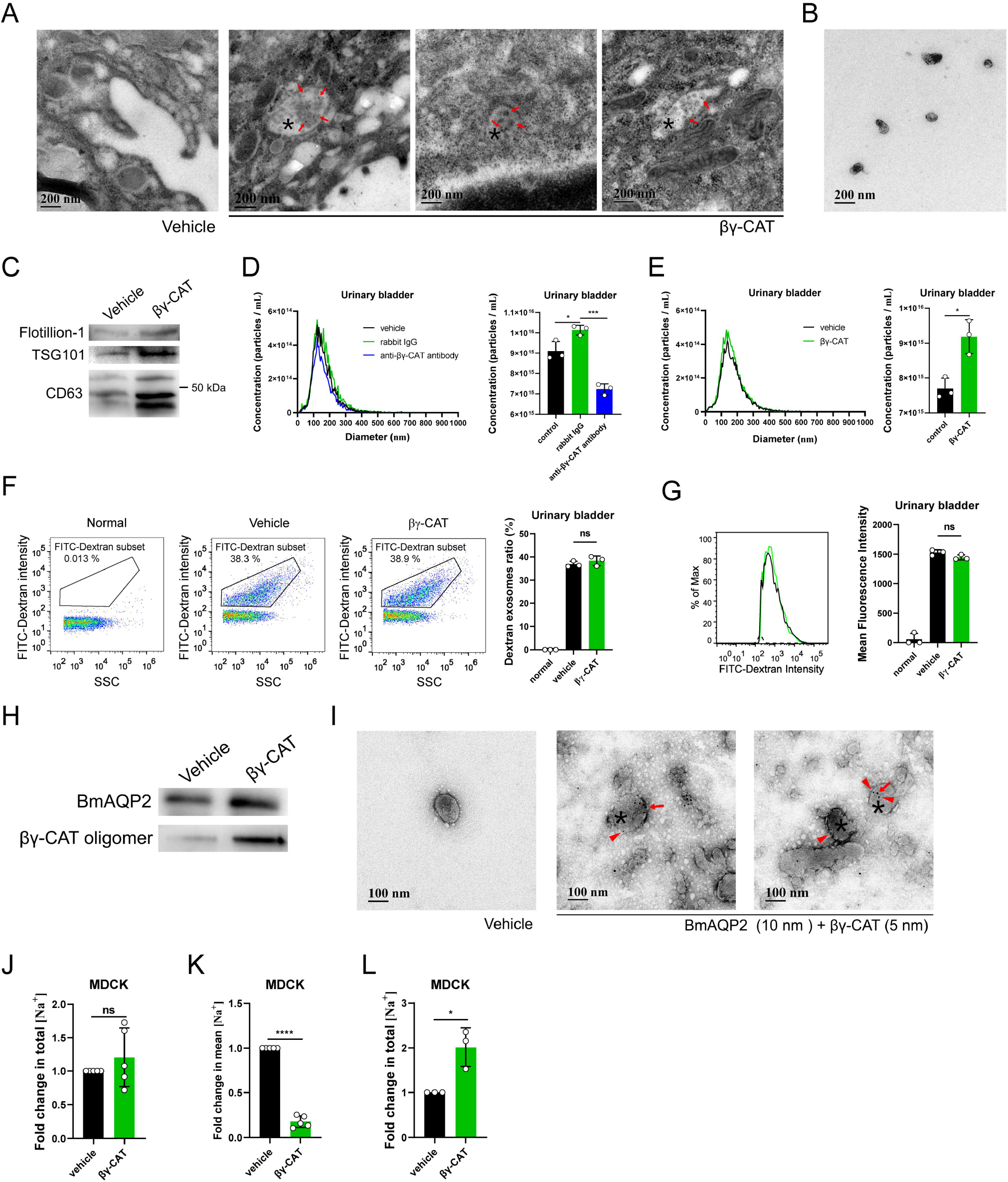
βγ-CAT enhances exosome release. (*A*) Ultrastructural localization of βγ-CAT in toad UB tissue by IEM. βγ-CAT (arrow) was readily detected in MVBs (asterisk) or ILVs. Vehicle (rabbit IgG control). (*B*) TEM analysis of exosomes from the supernatant of toad UB epithelial cells cultured *in vitro* for 3 hours. (*C*) Western blotting analysis of flotillin-1, TSG101 and CD63 molecules in exosomes isolated from the supernatant of toad UB epithelial cells cultured *in vitro* for 3 hours with or without the addition of 50 nM βγ-CAT. (*D, E*) Analysis of concentration and particle size (30-200 nm) of exosomes from toad UB epithelial cells with and without 50 μg/mL anti-βγ-CAT antibodies (*D*) or the addition of 50 nM βγ-CAT (*E*) by NTA. (*F, G*) The percentage (*F*) and mean fluorescence intensity (*G*) of dextran-containing exosomes from toad UB epithelial cells in a culture medium containing 1 mg/mL FITC-label dextran by Nanoflow Cytometry with or without the addition of 50 nM βγ-CAT. (*H*) Western blotting analysis of βγ-CAT and BmAQP2 in exosomes of toad UB epithelial cells cultured *in vitro* for 3 hours with or without the addition of 50 nM βγ-CAT. (*I*) IEM determination of βγ-CAT and BmAQP2 in exosomes (asterisk) from toad UB epithelial cells. BmAQP2 and βγ-CAT were labeled with 10-nm (arrow) and 5-nm (triangle) colloidal gold particles, respectively. (*J, K*) Fold changes of total (*J*) and mean (*K*) Na^+^ concentrations in exosomes from the same number of MDCK with or without 10 nM βγ-CAT for 3 hours. (*L*) Fold changes of total Na^+^ concentrations in exosomes from the same number of MDCK under the hypertonic medium with or without 10 nM βγ-CAT for 3 hours. The bars represent the mean ± SD of triplicate samples in *D*-*G*. Data represent the mean ± SD of at least three independent replicates in *J*-*L*. ns (*P* ≥ 0.05), \**P* < 0.05, \*\**P* < 0.01 and \*\*\*\**P* < 0.0001 by the unpaired *t* test. All data are representative of at least two independent experiments. See also Fig. S4.

We further investigated the properties of exosomes stimulated by βγ-CAT. The size distribution of exosomes derived from toad UB epithelial cells was determined by flow cytometry for nanoparticle analysis. The diameters of these exosomes were mainly concentrated in the range of 50–150 nm (Fig. S4*F*). When 1 mg/mL of FITC-dextran was incubated with isolated exosomes at room temperature for 3 hours, the exosomes did not contain FITC-dextran (Fig. S4*G*). In contrast, when dextran was added to the culture of toad UB epithelial cells under the same conditions, FITC-dextran was identified in exosomes secreted by the cells (Fig. S4*G*). This observation indicated that isolated exosomes did not take up 70-kDa dextran, but dextran was taken up by toad UB epithelial cells by macropinocytosis and released from the cells in the form of exosomes. Interestingly, the proportion of exosomes containing FITC-dextran remained unchanged in toad UB epithelial cells with or without the addition of purified βγ-CAT (Fig. 5*F*), and there was no difference in the mean fluorescence intensity of these exosomes (Fig. 5*G*). These results suggested that βγ-CAT promotes exocytosis by increasing exosome release, facilitating the transcellular transport of extracellular substances like dextran. However, the apparent properties of exosomes released seem not altered as assayed at the present stage.

Both BmAQP2 and βγ-CAT oligomers were detected by western blotting in exosomes released from toad UB epithelial cells, reflecting the presence of endogenous βγ-CAT. The addition of βγ-CAT to the cells greatly enhanced the quantity of BmAQP2 and βγ-CAT oligomers detected (Fig. 5*H*). IEM demonstrated colocalization of BmAQP2 and βγ-CAT in individual exosomes (Fig. 5*I*). These observations suggest that βγ-CAT drives the extracellular recycling and/or intercellular communication of AQPs via exosome release.

Furthermore, we analyzed the effect of βγ-CAT on Na^+^ levels in exosomes from MDCK cells. No difference was observed for total Na^+^ concentrations in exosomes produced by the same number of cells treated with or without 10 nM βγ-CAT (Fig. 5*J*). However, βγ-CAT increased exosome release of MDCK by 6.8 times (Fig. S4*D*), so the mean Na^+^ concentrations of exosomes were significantly reduced (Fig. 5*K*). In addition, an increase in total Na^+^ concentrations in exosomes induced by βγ-CAT were found in hypertonic media under the same conditions (Fig. 5*L*).

## Discussion

Amphibians live both on land and in water, and their skin is naked and acts as an important organ in water balance, respiration (gas exchange) and immune defense (1, 3). Though initially identified in *B. maxima* skin secretions (19), the PFP complex βγ-CAT is widely expressed in *B. maxima* water balance organs, including skin, UB and kidney (Fig 1 *C*-*H*, Fig. S1 *C* and *D*). Previous studies using RT-PCR likewise indicated the expression of βγ-CAT in various *B. maxima* tissues (23, 24). Interestingly, in contrast to its potent cytotoxicity to various mammalian cells (19, 25), βγ-CAT only shows toxic effects in toad cells in a dosage up to 2 μM (Fig. S2*B*), which is much higher than βγ-CAT physiological concentrations observed (23). Its toxicity to mammalian cells could reflect the absence of regulatory mechanisms for this exogenous PFP protein, which are normally present in the toad (18, 22). These observations support the notion that the PFP complex βγ-CAT can not be simply viewed as a cell death inducer and/or a microbicide, and besides its roles in immune defense (17, 23, 24), the protein exhibits other essential physiological functions in this toad.

*B. maxima* skin is covered with skin secretions containing various biological molecules that fulfill a range of physiological function, including hormone-like peptides, antimicrobial peptides, af-PFPs, TFFs, and haem b-containing albumin (16, 37, 38). Indeed, our results revealed that the skin secretions are necessary to prevent dehydration (Fig 1*B*). We further demonstrated that a major skin secretion component βγ-CAT has a role in toad water homeostasis. This PFP complex possesses the capacity to stimulate and participate in macropinocytosis (endocytosis) and exosome release (exocytosis) by epithelial cells of toad osmoregulatory organs, which promotes water and Na^+^ transport involved in water homeostasis.

Previous studies with mammalian cells and toad peritoneal cells showed that βγ-CAT exerts its biological actions by endocytosis and the formation of channels on endolysosomes (17, 19, 38), but the endocytic pathway of the protein remains elusive. Macropinocytosis, also referred to as cell drinking, is a form of endocytosis that mediates the non-selective uptake of extracellular fluid and solutes (6, 12). In a murine DC model, preliminary data showed that βγ-CAT increases the internalization of ovalbumin (antigen), presumed to be mediated by enhanced macropinocytosis (25). The present study carried out on various toad-derived cells, including epithelial cells from osmoregulatory organs and peritoneal cells, clearly showed that βγ-CAT is capable of inducing and participating in macropinocytosis. Macropinocytosis stimulated by βγ-CAT has been shown to facilitate the entry of water and Na^+^ into cells as assayed in epithelial cells from toad osmoregulatory organs (Fig. 2*F* and 3*F*). This may explain the role of βγ-CAT in promoting the recovery of cell volume under hypertonic stress (Fig. 2 *D*-*F*). Growth factor-induced macropinocytosis is mediated by the activation of the Ras and PI3-kinase signaling pathways (11). Similarly, activation of PI3-kinase and Akt signaling was involved in macropinocytosis stimulated by βγ-CAT, as suggested by the effects of pharmacological inhibitors and Akt phosphorylation analysis (Fig. 3 *G*-*I*). However, in contrast to classic growth factors, which bind to membrane protein receptors to initiate their signaling, βγ-CAT targets gangliosides and sulfatides as receptors in lipid rafts to initiate its cellular effects (22). The signals downstream from these lipid components, and their relationship to the induction of macropinocytosis, are presently unclear and are worthy of further study. In this respect, possible differences between macropinocytosis stimulated by growth factors and that induced by βγ-CAT should be investigated.

While engulfing large volumes of fluid, macropinocytosis also internalizes cell surface proteins such as receptors and integrins (39, 40). It has been proposed that endocytosis stimulated by βγ-CAT plays a role in the sorting of specific plasma membrane elements, such as functional integrated proteins or lipid components, which could help regulate cell responses to environmental variations (17). In addition, internalization and recycling of AQPs between the plasma membrane and the endosomal compartment have roles in controlling water uptake and conservation (5, 27). Accordingly, colocalization of βγ-CAT with AQPs was observed in endocytic organelles (Fig. 4), suggesting macropinocytosis induced by this PFP protein drives endocytosis and recycling of AQPs. Clearly, the presence of AQPs in the plasma membrane might facilitate water loss when cells faced by hypertonic osmotic stress, thus internalization of AQPs induced by βγ-CAT could counteract dehydration by preventing rapid water efflux and cell shrinkage.

The secretion of extracellular vesicles comprising exosomes and microvesicles represents a novel mode of intercellular communication and material exchange (36, 41). Previously, βγ-CAT was found to augment βγ-CAT-containing exosome release from murine DCs, which activates T cell response effectively (25). Because βγ-CAT is a factor exogenous to murine DCs, the explanation of the phenomenon is not obvious. However, the PFP is an endogenous element to toad cells. The present study illustrated that cell exocytosis in the form of exosome release was indeed augmented in the presence of βγ-CAT, as determined in diverse toad-derived cells (Fig. 5*D*, 5*E*, S4*B* and S4*C*), revealing that mediation of cell exocytosis via exosome release is an intrinsic property of this PFP protein. Previous studies demonstrated that βγ-CAT characteristically neutralizes the acidification of endocytic organelles containing this protein (23–25). This cellular process may result in the transformation of βγ-CAT-containing endocytic organelles into MVBs that does not fuse with lysosomes for the degradation of contained solutes. The observation that extracellular dextran, a tracer for βγ-CAT-induced endocytosis, was present in exosomes containing βγ-CAT (Fig. 5 *F* and *G*) supports this view.

The observation that βγ-CAT both stimulates and participates in macropinocytosis and exosome release strongly implies that this PFP is concerned with transcellular transport of internalized extracellular substances by PFP-driven vesicular transportation. This property is particularly important *in vivo*, where βγ-CAT could transport external substances such as water and Na^+^ to internal environments of toad tissues without disruption of the cellular tight junctions that maintain epithelial barrier functions. Interestingly, AQPs were identified in exosomes released in the presence of βγ-CAT (Fig 5 *H* and *I*), suggesting that βγ-CAT modulates water homeostasis by driving extracellular recycling of, and/or intercellular communication via these water channels. Alternatively, βγ-CAT-mediated exocytosis might function as a means for expulsion of noxious and indigestible solutes contained in fluid macropinocytosed by cells for water acquisition, like the expelling of dextran from toad UB epithelial cells (Fig. 5 *F* and *G*). The role of βγ-CAT in expelling noxious substances engulfed by macropinocytosis via exocytosis induction and the cellular sorting mechanism involved are important future challenges.

It is generally understood that classic membrane integrated proteins, like AQPs and ion channels play fundamental roles in water transport and homeostasis (27, 42). Furthermore, the tight junction protein claudin-2 mediates the permeability to transepithelial water movements under osmotic or Na^+^ gradients through the paracellular pathway (43–45). The present study has introduced a new player, a secretory endolysosome channel protein βγ-CAT, into the network of water transport and homeostasis via modulation of cellular endocytic and exocytic pathways. A cellular working model of the PFP βγ-CAT in promoting water uptake and maintaining was proposed in toad *B. maxima* (Fig. 6). In this model, water and Na^+^ can be taken up by macropinocytosis induced by the PFP βγ-CAT. Meanwhile, the transmembrane channels formed by βγ-CAT can mediate the Na^+^ flow (Fig 2 *B* and *C*). Previous studies have shown that the channel formed by βγ-CAT can cause cation ion flux (46). The PFP complex targets viral envelope to form pores that induced potassium and calcium ion efflux (21). Unexpectedly, the average Na^+^ concentrations of βγ-CAT-containing exosomes were substantially lower than those of the control (Fig 5 *J* and *K*). This phenomenon suggested that the presence of channels formed by βγ-CAT can lead to rapid efflux of Na^+^ out of endolysosomes and/or exosomes. Thus, Na^+^ can be release into cytosol through βγ-CAT channels in endocytic organelles, which drives water release to the cytosol via AQPs (Fig. 4). This is similar to the situation of two-pore channels working in mammalian macrophages (39, 40). Enhanced exosome release mediated by βγ-CAT may facilitate vesicular transport of ions such as Na^+^ (Fig. 5*L*), and/or water, into the intercellular spaces below tight junctions favoring water absorption into deeper cell environments. Meanwhile, membrane-integrated ion channels and AQPs in the basal plasma membrane of epithelial cells could mediate the efflux of Na^+^ and water into internal tissues underneath these cells achieving the absorption/reabsorption of water into toad internal tissues (5, 47). Drinking water by macropinocytosis is energetically expensive (12). However, the function of secretory PFP βγ-CAT in water acquisition and maintaining is necessary in light of various osmotic conditions that toad *B. maxima* has to face throughout its life cycle.

**Fig 6.**
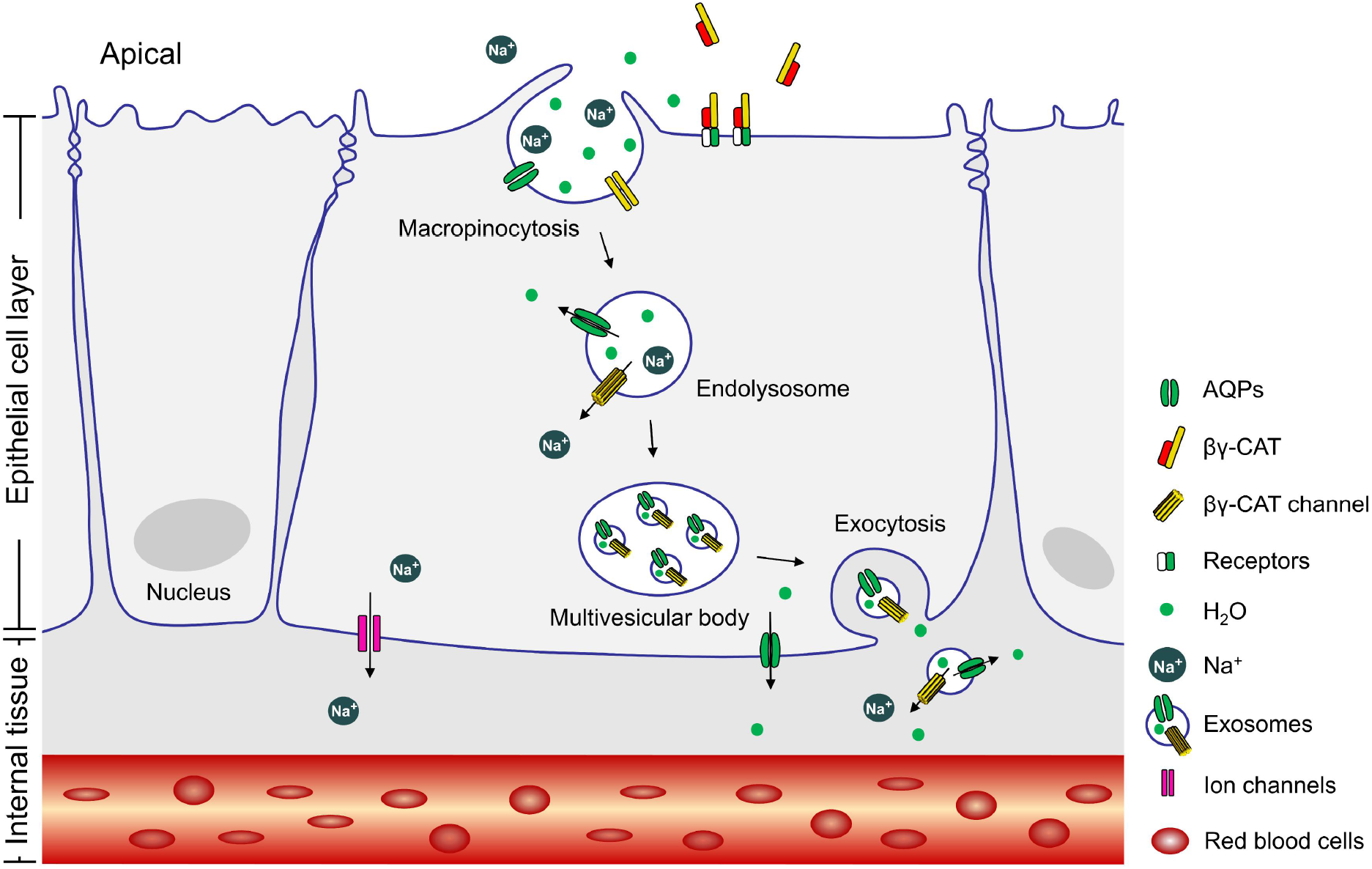
Proposed action model of βγ-CAT in water acquisition and maintaining. βγ-CAT achieved intracellular vesicle transport and transcellular transport of water, Na^+^ and AQP2 by promoting macropinocytosis (endocytosis) and exosome release (exocytosis). For a detailed description, see the test. Presentation of the epithelial cell layer and internal tissue is simplified.

Secretory PFPs have been identified in organisms from all kingdoms of life (15, 16), and af-PFPs are widely distributed in plants and animals (15–17). The present study provides functional evidence to support our previous hypothesis on the SELC (secretory endolysosome channel) pathway mediated via a secretory PFP in cellular endocytic and exocytic systems (17). Regulated relying on environmental cues (17, 18), SELC protein βγ-CAT is characterized of driving intracellular intake and transport of extracellular materials and/or plasma membrane components in a vesicle form. These capacities raise the possibility that besides working in water maintaining, βγ-CAT and/or its homologues could play active and fundamental roles in cell metabolism, as previously proposed (17). These SELC proteins and classic membrane integrated proteins (like solute carrier family members) can form two coordinated cell arms in extracellular nutrient sensing, sampling and acquiring. Furthermore, it is possible that βγ-CAT and its homologues are involved in modulating cellular metabolic responses to nutrient variation and shortage in toad *B. maxima*. Obviously, these SELC proteins should be essential to cells in those cases that classic membrane solute carriers are blockaded or even absent. Our findings will serve as clues for uncovering the physiological roles of PFPs in other living organisms. Specifically, proliferating embryonic cells of oviparous animals depend entirely on catabolism of extracellular macromolecules deposited in the egg white and yolk (10). If macropinocytosis-driven by a PFP plays a role in oviparous animal embryonic development is an interesting subject for further investigation.

In conclusion, the current work elucidated an unexpected role of a secretory PFP in water transport and homeostasis. βγ-CAT, an af-PFP and a TFF complex, induce and participate in macropinocytosis and exosome release in epithelial cells from *B. maxima* osmoregulatory organs. These effects could explain the role of the PFP complex in promoting the internalization, transport and release of water, Na^+^ and AQPs, working together in the maintenance of toad cell volume regulation and water homeostasis (Fig. 6). Together with membrane-integrated AQPs and ion channels, the action of secretory βγ-CAT can promote the overall achievement of water maintaining and balance, especially when the toad encounters osmotic stress.

## Materials and Methods

### Animals

Toads (*B. maxima*) were captured in the wild and raised at room temperature by feeding with live *Tenebrio molitor*. Toads with an average weight of 19 ± 5 g were used in experiments after fasting in isotonic Ringer’s solution for 3 days. All the procedures and the care and handling of the animals were approved by the Institutional Animal Care and Use Committee at Kunming Institute of Zoology, Chinese Academy of Sciences (Approval ID: IACUC-OE-2021-05-001).

### Animal experiment

Formulation of isotonic Ringer’s solution (111.2 mM NaCl, 1.9 mM KCl, 1.1 mM CaCl_2_, 2.4 mM NaHCO_3_, 1.6 mM MgCl_2_) was modified based on a previous report (48). After fasting, toads in the hypertonic group were placed in hypertonic Ringer’s solution (Ringer’s solution with 222.4 mM NaCl) for 3 hours and their weight changes were recorded. Toads in the hypertonic/isotonic group were then transferred from hypertonic Ringer’s solution to isotonic Ringer’s solution for 3 hours and their weight changes were recorded. Weight changes of toads in isotonic Ringer’s solution were recorded for 6 hours as a reference, termed the isotonic group. The skin of some toads was electrically stimulated (4.5 V DC, pulse duration 10 ms) for 3 minutes to deplete skin secretions, and the animals were placed in isotonic Ringer’s solution for 30 minutes. Their survival rates were then recorded after 48 hours in isotonic and hypertonic solutions.

### Cell culture

Toad epithelial cells were obtained by tissue digestion. Specifically, the UB, skin, and kidney tissue were dissected from five toads whose spinal cord had been destroyed. Tissues were further rinsed and stripped in Ringer’s solution to remove residual blood, mucus and other impurities. They were then cut into pieces and washed twice in Ringer’s solution, followed by oscillatory digestion with trypsin at room temperature for 40 minutes. The cells were separated with a 200-mesh sieve. In addition, toad peritoneal cells were extracted from the peritoneal fluid of *B. maxima*. All cells were centrifugally enriched with 2,000 rpm for 5 min at 4℃. The digestive cells in toad skin and UB tissues contain around 90% and 60% of epidermal cells, respectively, by flow cytometry using an anti-pan cytokeratin AE1/AE3 monoclonal antibody (Thermo Fisher Scientific, Rockford, IL, USA).

### Additional Assays

All other materials and methods are described in Materials and Methods of Supplemental Information.

## Acknowledgements

This work was supported by grants from the National Natural Science Foundation of China (grant numbers 31872226, U1602225 and 31572268) and the Yunling Scholar Program to Yun Zhang. We would like to thank the Kunming Biological Diversity Regional Center of Instrument, Kunming Institute of Zoology, Chinese Academy of Sciences for our electron microscopy and we would be grateful to Yingqi Guo for her help of making EM samples.

## Author contributions

Y.Z., Z.Z., and Z.H.S conceived of and conceptualized the study; Z.Z., Z.H.S., and C.J.Y did the experiments, Z.Z., Y.Z., Z.H.S, and C.J.Y analyzed and interpreted the data; Z.Z., Y.Z., Z.H.S wrote the manuscript; Y.Z., Z.Z., Z.H.S and C.J.Y critically revised the manuscript for important intellectual content.

## References

1. P. J. Bentley, Adaptations of amphibia to arid environments. Science 152, 619–623 (1966).

2. V. Shoemaker, K. A. Nagy, Osmoregulation in amphibians and reptiles. Annu. Rev. Physiol. 39, 449–471 (1977).

3. S. D. Hillyard, Behavioral, molecular and integrative mechanisms of amphibian osmoregulation. J. Exp. Zool. 283, 662–674 (1999).

4. Y. Ogushi, et al., The water-absorption region of ventral skin of several semiterrestrial and aquatic anuran amphibians identified by aquaporins. Am. J. Physiol.-Regul. Integr. Comp. Physiol. 299, R1150–R1162 (2010).

5. M. Suzuki, T. Hasegawa, Y. Ogushi, S. Tanaka, Amphibian aquaporins and adaptation to terrestrial environments: A review. Comp. Biochem. Physiol. A. Mol. Integr. Physiol. 148, 72–81 (2007).

6. M. C. Kerr, R. D. Teasdale, Defining macropinocytosis. Traffic 10, 364–371 (2009).

7. J. A. Swanson, C. Watts, Macropinocytosis. Trends Cell Biol. 5, 424–428 (1995).

8. C. Ramirez, A. D. Hauser, E. A. Vucic, D. Bar-Sagi, Plasma membrane V-ATPase controls oncogenic RAS-induced macropinocytosis. Nature 576, 477–481 (2019).

9. M. Bretou, et al., Lysosome signaling controls the migration of dendritic cells. Sci. Immunol. 2 (2017).

10. W. Palm, C. B. Thompson, Nutrient acquisition strategies of mammalian cells. Nature 546, 234–242 (2017).

11. W. Palm, Metabolic functions of macropinocytosis. Philos. Trans. R. Soc. B Biol. Sci. 374, 20180285 (2019).

12. J. A. Swanson, J. S. King, The breadth of macropinocytosis research. Philos. Trans. R. Soc. B Biol. Sci. 374, 20180146 (2019).

13. M. W. Parker, et al., Structure of the Aeromonas toxin proaerolysin in its water-soluble and membrane-channel states. Nature 367, 292–295 (1994).

14. M. D. Peraro, F. G. van der Goot, Pore-forming toxins: ancient, but never really out of fashion. Nat. Rev. Microbiol. 14, 77–92 (2016).

15. P. Szczesny, et al., Extending the aerolysin family: from bacteria to vertebrates. PLoS ONE 6, e20349 (2011).

16. Y. Zhang, Why do we study animal toxins? Zool. Res. 36, 183–222 (2015).

17. Y. Zhang, Q. Q. Wang, Z. Zhao, C. J. Deng, Animal secretory endolysosome channel discovery. Zool. Res. 42, 141–152 (2021).

18. Q. Q. Wang, et al., A cellular endolysosome-modulating pore-forming protein from a toad is negatively regulated by its paralog under oxidizing conditions. J. Biol. Chem. 295, 10293–10306 (2020).

19. S. B. Liu, et al., A novel non-lens βγ-crystallin and trefoil factor complex from amphibian skin and its functional implications. PLoS ONE 3, e1770 (2008).

20. Q. Gao, et al., Characterization of the βγ-crystallin domains of βγ-CAT, a non-lens βγ-crystallin and trefoil factor complex, from the skin of the toad Bombina maxima. Biochimie 93, 1865–1872 (2011).

21. L. Liu, et al., An aerolysin-like pore-forming protein complex targets viral envelope to inactivate Herpes Simplex Virus Type 1. J. Immunol. (2021).

22. X. L. Guo, et al., Endogenous pore-forming protein complex targets acidic glycosphingolipids in lipid rafts to initiate endolysosome regulation. Commun. Biol. 2, 59 (2019).

23. Y. Xiang, et al., Host-derived, pore-forming toxin-like protein and trefoil factor complex protects the host against microbial infection. Proc. Natl. Acad. Sci. 111, 6702–6707 (2014).

24. S. A. Li, et al., Host pore-forming protein complex neutralizes the acidification of endocytic organelles to counteract intracellular pathogens. J. Infect. Dis. 215, 1753–1763 (2017).

25. C. J. Deng, et al., A secreted pore-forming protein modulates cellular endolysosomes to augment antigen presentation. FASEB J. 34, 13609–13625 (2020).

26. Z. H. Gao, et al., Pore-forming toxin-like protein complex expressed by frog promotes tissue repair. FASEB J. 33, 782–795 (2019).

27. K. Takata, Aquaporins: water channel proteins of the cell membrane. Prog. Histochem. Cytochem. 39, 1–83 (2004).

28. A. de Baey, A. Lanzavecchia, The role of aquaporins in dendritic cell macropinocytosis. J. Exp. Med. 191, 743–748 (2000).

29. M. Nonnenmacher, T. Weber, Adeno-associated virus 2 infection requires endocytosis through the CLIC/GEEC pathway. Cell Host Microbe 10, 563–576 (2011).

30. J. P. Lim, P. A. Gleeson, Macropinocytosis: an endocytic pathway for internalising large gulps. Immunol. Cell Biol. 89, 836–843 (2011).

31. N. Araki, M. T. Johnson, J. A. Swanson, A role for phosphoinositide 3-kinase in the completion of macropinocytosis and phagocytosis by macrophages. J. Cell Biol. 135, 1249–1260 (1996).

32. H. P. Lin, et al., Identification of novel macropinocytosis inhibitors using a rational screen of Food and Drug Administration-approved drugs. Br. J. Pharmacol. 175, 3640–3655 (2018).

33. R. M. Hays, A. Leaf, Studies on the movement of water through the isolated toad bladder and its modification by vasopressin. J. Gen. Physiol. 45, 905–919 (1962).

34. T. Hasegawa, M. Suzuki, S. Tanaka, Immunocytochemical studies on translocation of phosphorylated aquaporin-h2 protein in granular cells of the frog urinary bladder before and after stimulation with vasotocin. Cell Tissue Res. 322, 407–415 (2005).

35. C. W. Davis, A. L. Finn, Interactions of sodium transport, cell volume, and calcium in frog urinary bladder. J. Gen. Physiol. 89, 687–702 (1987).

36. G. van Niel, G. D’Angelo, G. Raposo, Shedding light on the cell biology of extracellular vesicles. Nat. Rev. Mol. Cell Biol. 19, 213–228 (2018).

37. Y. X. Zhang, R. Lai, W. H. Lee, Y. Zhang, Frog albumin is expressed in skin and characterized as a novel potent trypsin inhibitor. Protein Sci. 14, 2469–2477 (2005).

38. F. Zhao, et al., Comprehensive transcriptome profiling and functional analysis of the frog (Bombina maxima) immune system. DNA Res. 21, 1–13 (2014).

39. S. A. Freeman, et al., Lipid-gated monovalent ion fluxes regulate endocytic traffic and support immune surveillance. Science 367, 301–305 (2020).

40. J. S. King, E. Smythe, Water loss regulates cell and vesicle volume. Science 367, 246–247 (2020).

41. A. E. Russell, et al., Biological membranes in EV biogenesis, stability, uptake, and cargo transfer: an ISEV position paper arising from the ISEV membranes and EVs workshop. J. Extracell. Vesicles 8, 1684862 (2019).

42. M. L. McManus, K. B. Churchwell, K. Strange, Regulation of cell volume in health and disease. N. Engl. J. Med. 333, 1260–1267 (1995).

43. R. Rosenthal, et al., Claudin-2, a component of the tight junction, forms a paracellular water channel. J. Cell Sci. 123, 1913–1921 (2010).

44. A. Wilmes, L. Aschauer, A. Limonciel, W. Pfaller, P. Jennings, Evidence for a role of claudin 2 as a proximal tubular stress responsive paracellular water channel. Toxicol. Appl. Pharmacol. 279, 163–172 (2014).

45. R. Rosenthal, et al., Claudin-2-mediated cation and water transport share a common pore. Acta Physiol. 219, 521–536 (2017).

46. Q. Gao, et al., βγ-CAT, a non-lens betagamma-crystallin and trefoil factor complex, induces calcium-dependent platelet apoptosis. Thromb. Haemost. 105, 846–854 (2011).

47. Y. Marunaka, Characteristics and pharmacological regulation of epithelial Na+ channel (ENaC) and epithelial Na+ transport. J. Pharmacol. Sci. 126, 21–36 (2014).

48. A. Edström., Effects of Ca2+ and Mg2+ on rapid axonal transport of proteins in vitro in frog sciatic nerves. J. Cell Biol. 61, 812–818 (1974).

