## Supplemental Information for "A pore-forming protein drives macropinocytosis to facilitate toad water maintaining"

### Materials and Methods

#### Cell culture

Madin-Darby canine kidney (MDCK), Caco-2, HEK293 and T24 cell lines were purchased from Kunming Cell Bank, Chinese Academy of Sciences. Cells were cultured in DMEM/F-12 (Biological Industries, Cat 01-172-1A) containing 10% fetal bovine serum (Biological Industries, Cat 04-001-1A) and 1% levofloxacin hydrochloride and sodium chloride injection.

#### Purification of βγ-CAT

βγ-CAT purification and purity analysis were carried out according to the previous description (1, 2).

#### Cell viability assay

The MTS assay was used to detect the cytotoxicity of βγ-CAT. In brief, MDCK and T24 cell lines were seeded in 96-well plates at $1\times10^4$ cells per well and cultured overnight at 37°C in 5% $CO_2$. Cells were incubated with the MTS reagent (Promega, Cat G3580) in the dark for 1 hour after treatment with βγ-CAT at room temperature for 2 hours. The absorbance of the toad cells culture supernatant was measured with $2\times10^6$ cells per well. The absorption was detected at 490 nm with the Infinite 200 Pro microplate reader (Tecan, Männedorf, Switzerland).

#### Hemolytic activity

$2\times10^7$ toad skin, UB, kidney or peritoneal cells were centrifugally enriched and then washed three times with 5 mL Ringer's solution until the supernatant had no hemolytic activity on human erythrocytes. The toad cells suspension was resuspended into four groups. The uncultured control group was centrifuged and the supernatant was collected. The other three groups were incubated at room temperature for 1.5 hours and centrifuged to collect the supernatant. One of them served as the cultured group. The remaining two batches were mixed with 50 μg/mL rabbit antibody or rabbit-derived anti-βγ-CAT antibody, respectively, and incubated at room temperature for 30 minutes, while the uncultured and cultured groups were treated similarly using the same volume of Ringer's solution. Human erythrocytes were added to the supernatant at a concentration of 3%, incubated for 30 min at 37 ℃, and then centrifuged at 2,000 rpm for 5 min. The absorption was detected at 540 nm with the Infinite 200 Pro microplate reader (Tecan, Männedorf, Switzerland).

**Sequence alignment and sequence analysis**

The evolutionary history was inferred by using the Maximum Likelihood method based on the Poisson correction model (3). Initial tree(s) for the heuristic search were obtained automatically by applying Neighbor-Join and BioNJ algorithms to a matrix of pairwise distances estimated using a JTT model, and then selecting the topology with superior log likelihood value (4). Evolutionary analyses were conducted in MEGA7. In addition, sequence alignment analysis of AQPs in *B. maxima* and other species was done by Clustal Omega.

**Cell diameter measurement**

Previously described methods were modified as appropriate (5). Briefly, digested MDCK, Caco-2 and T24 cells were cultured in serum-free medium for 30 minutes at a concentration of $2\times10^6$ cells/mL before detection. To determine cell diameters, we treated MDCK, Caco-2 and T24 cells with hypertonic PBS (adding NaCl to 0.01 M PBS increased osmotic pressure to 400 mOsm) containing purified βγ-CAT at doses of 10 nM, 10 nM and 5 nM, respectively. Toad UB epithelial cells were incubated with or without 0.3 mM $HgCl_2$ for 10 minutes at a concentration of $2\times10^6$ cells/mL after incubating with 50 μg/mL rabbit-derived anti-βγ-CAT antibody for 30 minutes at room temperature. Diameter changes of toad UB epithelial cells were detected in isotonic or hypertonic Ringer's solution. The final concentration of all cells was $1\times10^6$ cells/mL. The average cell diameter was determined by a Countstar Automated Cell Counter (ALIT Life Science, Shanghai, China).

### Patch-clamp recordings

Currents were recorded on HEK-293 cells with the outside-out configuration of the patch-clamp technique at room temperature with a Multiclamp700B amplifier and a Digidata 1550A analog-digital converter controlled by a pClamp10 software (Molecular Devices, San Jose, USA). The pipette solution contained 150 mM KCl, 10 mM HEPES, 1 mM EGTA, pH7.4; the bath solution contained 150 mM NaCl, 10 mM HEPES, pH7.4. The background (blank) and macroscopic current induced by 100 nM βγ-CAT were recorded under 500 ms ramp protocol between −100 mV to +100 mV every 2 s from a holding potential of 0 mV. For on replacement experiments, the pipette solution contained 150 mM NaCl, 10 mM HEPES, 1 mM EGTA, pH7.4; the bath solution contained 150 mM or 15 mM NaCl, 10 mM HEPES, pH7.4. All currents shown have been leak subtracted. Data were analyzed with pClamp10 and GraphPad Prism 8.

### Quantitative real-time PCR

The mRNA levels of βγ-CAT-α and βγ-CAT-β in skin, UB and kidney were detected by qRT-PCR using a Hieff qPCR SYBR Green Master Mix (No Rox) kit (Yeasen, Cat 11201ES03). The cycle counts of target genes were normalized to those of β-actin. Primer sequences in this study are shown in **Table S1**.

### Immunofluorescence and HE staining

Appropriate adjustments were used on a previous description (6). Briefly, MDCK cells were grown to 90% on a 24-well glass slide. MDCK and UB epidermal cells were treated with a dose of 10 nM or 50 nM βγ-CAT, respectively, in 0.01 M PBS with or without 1 mg/mL FITC-label dextran (Sigma, Cat 46945) for 15 minutes at 37℃. After washing with PBS for three times, cells were fixed in 4% paraformaldehyde for 15 minutes, and then treated with 0.5% TritonX-100 for 15 minutes. After blocking for 2 hours in PBS containing 2% BSA at 37℃, cells were incubated with mouse-derived anti-AQP2 (Santa Cruz, Cat sc-515798), mouse-derived anti-AQP3 (Santa Cruz, Cat sc-518001) or rabbit-derived anti-βγ-CAT primary antibody for 2 hours at 37°C. Cells were incubated in the dark with fluorescence labeled secondary antibody for 1 hour at 37°C after washing with PBS for three times. The samples were sealed with an anti-fluorescent quench agent containing DAPI. Images were acquired by a Nikon A1 confocal laser microscope system (Nikon, Tokyo, Japan). The localization of βγ-CAT or BmAQP2 in toad tissues were determined by using rabbit-derived anti-AQP2 (ImmunoWay, Cat YT0290) and mouse-derived anti-βγ-CAT primary antibodies. Images were acquired by a Zeiss LSM 880 microscope system (Carl Zeiss, Oberkochen, Germany).

**Isolation of exosomes**

The method to isolate exosomes was optimized based on a previous description (7). Briefly, MDCK and T24 cells were grown to 90% on 100-mm cell culture dishes. The cells were cultured in twenty cell culture dishes with and without 10 nM or 5 nM βγ-CAT for 3 hours at 37°C. Cell supernatant was collected for gradient centrifugation (300 ×*g* for 30 minutes; 500 ×*g* for 30 minutes; 2,000 ×*g* for 30 minutes; 10,000 ×*g* for 40 minutes) at 4°C to remove residual cells, debris, and microvesicles. Exosomes were obtained by ultracentrifugation at 100,000 ×*g* for 2 hours at 4°C, and then resuspended in PBS, and enriched again in the CP100WX preparative ultracentrifuge (HATACHI, Tokyo, Japan).

$2\times10^7$ toad cells were cultured in Ringer's solution containing either 50 nM βγ-CAT, 50 μg/mL rabbit IgG or 50 μg/mL rabbit-derived anti-βγ-CAT antibodies for 3 hours at room temperature. Exosomes were extracted with Hieff Quick exosome isolation kit (Yeasen, Cat 41201-A). Cell supernatant was obtained by a series of gradient centrifugations (500 ×*g* for 10 minutes; 3,000 ×*g* for 10 minutes). It was thoroughly mixed with a quarter volume of the reagent at 4°C for 2 hours and then centrifuged at 10,000 ×*g* for 1 hour to collect exosomes. After resuspension in Ringer's solution, the liquid containing exosomes was centrifuged at 100,000 ×*g* for 2 hours at 4°C, and repeated once in a CP100WX preparative ultracentrifuge (HATACHI, Tokyo, Japan).

**Flow cytometry**

$2\times10^5$ MDCK and T24 cells were incubated with 100 μg/mL 70 kDa FITC-label dextran (Sigma, Cat 46945) or Lucifer Yellow (Sigma, Cat L0144) in the dark at 37°C for 30 minutes with and without 10 nM or 5 nM βγ-CAT, respectively. The samples were followed by fluorescence detection of FITC or AmCyan. In each sample, $1\times10^4$ single cells were analyzed. $2\times10^6$ toad UB epithelial cells and peritoneal cells were treated with 50 nM βγ-CAT, while 100 nM βγ-CAT was used for $2\times10^6$ toad skin and kidney cells. In the test using immunodepletion of endogenous βγ-CAT, toad cells were incubated with 50 μg/mL rabbit-derived anti-βγ-CAT antibodies for 30 minutes before the above protocol was carried out. During the inhibitor experiment, cells were first incubated with 100 μM EIPA (MedChemExpress, Cat HY-101840A) or 20 μM wortmannin (Sigma, Cat 681675) for 1 hour at 37°C. In addition, $2\times10^6$ digested toad UB epithelial cells were cultured *in vitro* for 3 hours and co-incubated with 500 ng/mL propidium iodide (Becton Dickinson, Franklin Lakes, NJ, USA) for 10 minutes at room temperature. The fluorescence was recorded using LSR Fortessa cell analyzer (Becton Dickinson, Franklin Lakes, NJ, USA). The data were analyzed by FlowJo 10 and GraphPad Prism 8.

**Flow cytometry for nanoparticle analysis**

Exosomes of the toad UB epithelial cells were isolated as described in "**Isolation of exosomes**". Exosomes or $2\times10^7$ toad UB epithelial cells were cultured in medium containing 1 mg/mL FITC-label dextran with or without 50 nM βγ-CAT at room temperature for 3 hours. Exosomes of the toad UB epithelial cells were enriched and residual dextran was washed off using the method described in "**Isolation of exosomes**". All experiments were done in the dark. The fluorescence and particle size distribution were recorded using Flow NanoAnalyzer (NanoFCM, Xiamen, China). The data were analyzed by FlowJo 10 and GraphPad Prism 8.

**Transmission electron microscopy (TEM) and Immunoelectron microscopy (IEM)**

The experiments were based on the previous report (8). Tissue samples were fixed overnight with 0.1 M PB (pH 7.4) containing 3% paraformaldehyde 0.1% glutaraldehyde at 4℃, washed with 0.1 M PB four times for 15 minutes, and then washed with 0.1 M glycine in 0.1 M PB for 30 minutes at 4℃. After ethanol gradient dehydration, the sample was embedded in LR white resin (sigma, Cat L9774) and polymerized at 55℃ for 24 hours. 100-nm ultrathin sections were prepared using an EM UC7 ultramicrotome (Leica Microsystems, Wetzlar, Germany) and loaded onto 200-mesh Ni grids (EMCN, Cat BZ10262Na). Enriched exosomes were directly attached to Ni grids for 10 minutes before experimental treatment. The samples were washed with deionized water for 2 minutes and then blocked with 1% BSA for 5 minutes. After overnight incubation with mouse-derived anti-βγ-CAT or rabbit-derived anti-AQP2 (Immunoway, Cat YT0290) primary antibodies at 4℃, the samples were washed with deionized water 10 times. Then, the samples were incubated with 5-nm or 10-nm colloidal gold-conjugated secondary antibody (sigma, Cat G7527 and G7402) at room temperature for 2 hours, and washed 10 times at room temperature. Exosomes were stained with 2% uranyl acetate for 3 minutes, and sections were stained with 2% uranyl acetate for 7 minutes and lead citrate for 5 minutes before observation by employing a JEM 1400 plus transmission electron microscope at 100 kV.

**Intracellular and exosome sodium detection**

$2\times10^7$ toad UB epithelial cells enriched by centrifugation were resuspended with Ringer's solution (N-Methyl-D-glucamine was used instead of NaCl) after digestion and washing 3 times. MDCK was cultured to 90% on 100-mm cell culture dishes, and three cell culture dishes in each group were used in this experiment. Toad UB epithelial cells and MDCK were cultured for 3 hours with and without 50 nM or 10 nM βγ-CAT, respectively. Cells were collected, and lysed with deionized water.

Sodium concentrations in MDCK exosomes were measured in isotonic or hypertonic assays using forty or twenty cell culture dishes in each group, respectively. MDCK exosomes were isolated as described in "**Isolation of exosomes**". The samples were frozen in liquid nitrogen and then slowly melted at room temperature, and repeated twice. Ultrasonic cell crushing (power 200W, work every 10 seconds for 5 seconds, 40 cycles) was used to release the sample until the liquid was transparent. Impurities in samples were removed by gradient centrifugation (2,000 ×*g* for 30 minutes; 10,000 ×*g* for 1 hour). After the sample was filtered with a 0.22 μm filter, the concentration of sodium ions in the samples was determined by iCAP6300 inductively coupled plasma optical emission spectrometer (Thermo Fisher Scientific, Waltham, United States of America).

**Nanoparticle Tracking Analysis (NTA)**

All exosome samples were diluted to around $1 \times 10^7$ particles/mL with PBS. The particle size and concentration of exosomes were measured by ZetaView PMX 110 (Particle Metrix, Meerbusch, Germany), and the data were analyzed using the software ZetaView 8.04.02.

**Western Blotting**

To measure the protein level of βγ-CAT or AQP2, toad UB epithelial cells, MDCK, Caco-2 and T24 cells were treated for 15 minutes with or without 50 nM, 10 nM, 10 nM and 5 nM βγ-CAT, respectively. Total Akt and phospho-Akt were detected by treating MDCK cells with or without 10 nM βγ-CAT for 30 minutes or 60 minutes. Exosomes from toad UB epithelial cells were washed and lysed for Western blotting. The rabbit-derived anti-βγ-CAT, rabbit-derived anti-AQP2 (Immunoway, Cat YT0290), rabbit-derived anti-total Akt (Solarbio, Cat K106557P) and rabbit-derived anti-phospho-Akt-T308 (Solarbio, Cat K006214P) primary antibodies were used in these experiments.

### Statistical analysis

Animal survival data were analyzed by the Gehan-Breslow-Wilcoxon test. All other data were analyzed using the standard unpaired t-test. Differences with $P$ values < 0.05 were considered statistically significance. All statistical analyses were conducted using GraphPad Prism 8 software.

### References


1. S. B. Liu, *et al.*, A novel non-lens βγ−crystallin and trefoil factor complex from amphibian skin and its functional implications. *PLoS ONE* **3**, e1770 (2008).

2. Y. Xiang, *et al.*, Host-derived, pore-forming toxin-like protein and trefoil factor complex protects the host against microbial infection. *Proc. Natl. Acad. Sci.* **111**, 6702–6707 (2014).

3. E. Zuckerkandl, L. Pauling, "Evolutionary Divergence and Convergence in Proteins" in *Evolving Genes and Proteins*, (Elsevier, 1965), pp. 97–166.

4. V. Hollich, L. Milchert, L. Arvestad, E. L. L. Sonnhammer, Assessment of Protein Distance Measures and Tree-Building Methods for Phylogenetic Tree Reconstruction. *Mol. Biol. Evol.* **22**, 2257–2264 (2005).

5. T. Marchbank, R. J. Playford, Trefoil factor family peptides enhance cell migration by increasing cellular osmotic permeability and aquaporin 3 levels. *FASEB J.* **32**, 1017–1024 (2018).

6. L. Li, H. Zhang, M. Varrin-Doyer, S. S. Zamvil, A. S. Verkman, Proinflammatory role of aquaporin-4 in autoimmune neuroinflammation. *FASEB J.* **25**, 1556–1566 (2011).

7. C. J. Deng, *et al.*, A secreted pore-forming protein modulates cellular endolysosomes to augment antigen presentation. *FASEB J.* **34**, 13609–13625 (2020).

8. J. Wang, *et al.*, GPRC5A suppresses protein synthesis at the endoplasmic reticulum to prevent radiation-induced lung tumorigenesis. *Nat. Commun.* **7**, 11795 (2016).

### Data and Figures

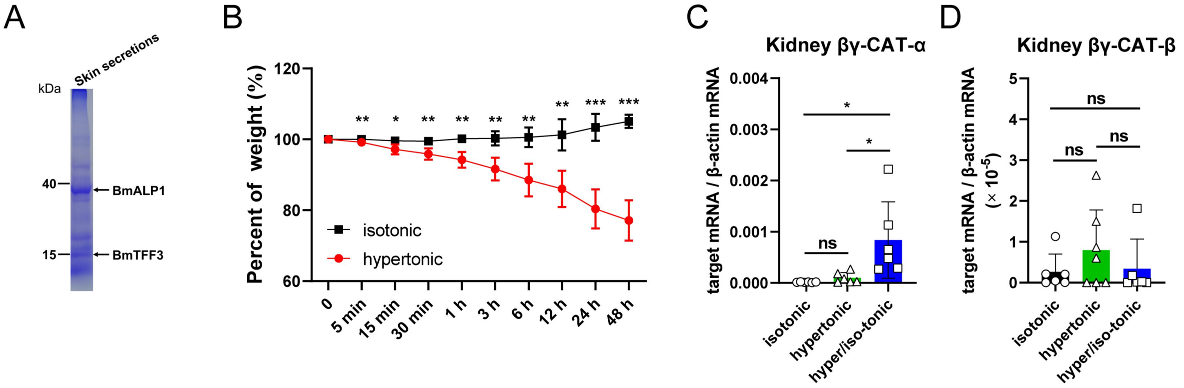


**Fig. S1. βγ-CAT is involved in responses to osmotic stress.** (*A*) 5 μg freeze-dried skin secretions were analyzed by SDS-PAGE with Coomassie blue staining. (*B*) Percentage change curves of toad weight in isotonic or hypertonic Ringer's solution within 48 hours ($n = 4$). (*C*, *D*) The expression of βγ-CAT subunits in toad kidney as analyzed by real-time fluorescent quantitative PCR after the animals were placed in isotonic, hypertonic and hypertonic/isotonic Ringer's solution (transferring toads from hypertonic to isotonic Ringer's solution) for 3 hours ($n = 6$). All data of the weight change and gene expression represent the mean ± SD. ns ($P \geq 0.05$), *$P < 0.05$, **$P < 0.01$ and ***$P < 0.001$ by unpaired $t$ test. All data are representative of at least two independent experiments.

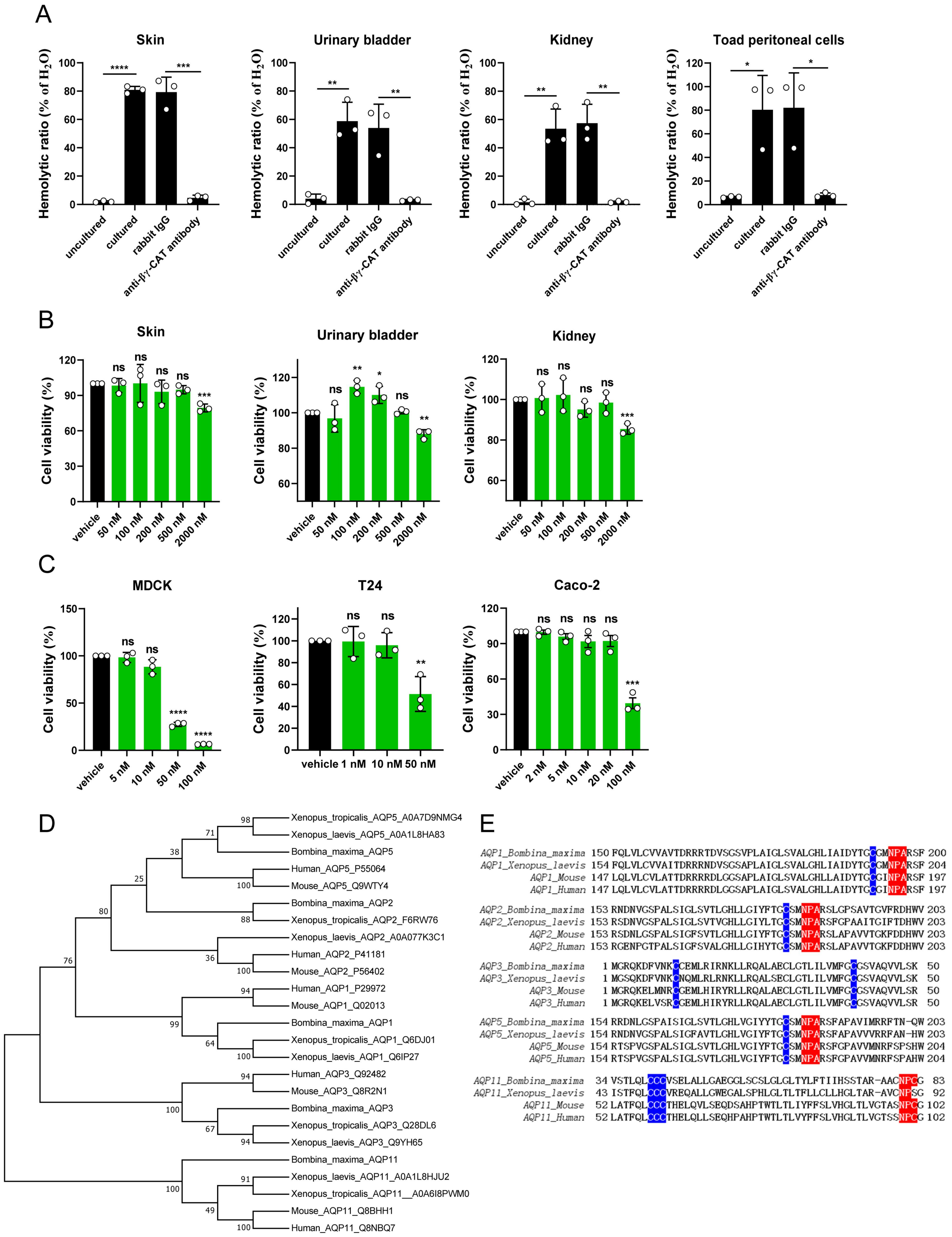


**Fig. S2. The endogenous secretion and cytotoxicity of βγ-CAT and toad *B. maxima* UB AQP analysis.** (*A*) The endogenous secretion of βγ-CAT was analyzed by its hemolytic activity on human erythrocytes. The samples assayed were uncultured supernatant, cultured supernatant, and cultured supernatant containing 50 μg/mL rabbit IgG supernatant or 50 μg/mL anti-βγ-CAT antibody supernatant for toad skin, UB, kidney and peritoneal cells. (*B*) Cytotoxicity assays of βγ-CAT on digestive epithelial cells of toad skin, UB and kidney. The cells were treated by βγ-CAT for 3 hours, then the cell viability was determined by MTS. (*C*) Cytotoxicity assays of βγ-CAT on MDCK, Caco-2 and T24 cells as assayed in (*B*). (*D*) Molecular phylogenetic analysis of AQPs in *B. maxima* UB and other species by Maximum Likelihood method. (*E*) Sequence alignment of AQPs in *B. maxima* UB and other species by Clustal Omega. NPA/NPC (red) is the signature motif of AQPs, and cysteine (blue) is the mercury binding site. The results are reported as mean ± SD of triplicate samples in *B*, *C*. ns ($P \geq 0.05$), *$P < 0.05$, **$P < 0.01$, ***$P < 0.001$ and ****$P < 0.0001$ by unpaired *t* test. All data are representative of at least two independent experiments in *A*-*C*.

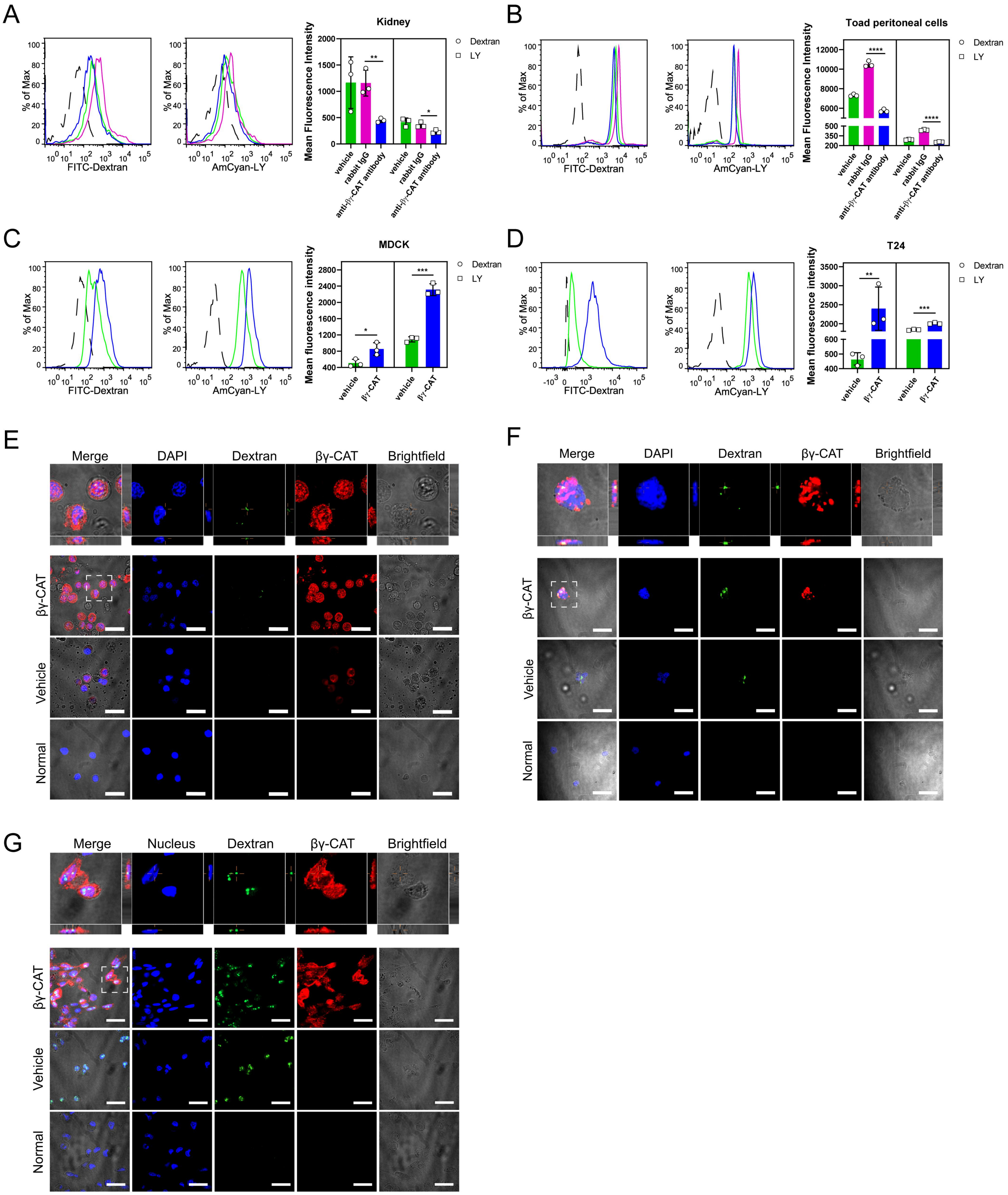


**Fig. S3. βγ-CAT promotes macropinocytosis.** (*A*, *B*) Immunedepletion of endogenous βγ-CAT decreased macropinocytosis. The mean fluorescence intensity of 70 kDa FITC-label dextran and Lucifer Yellow (LY) in toad kidney epithelial cells (*A*)

and peritoneal cells (*B*) was determined by flow cytometry. The cells were incubated with 50 µg/mL anti-βγ-CAT antibodies to immunodeplete endogenous βγ-CAT for 30 minutes, and rabbit IgG was used as an antibody control. Vehicle refers to an antibody absent control. Then the cells were incubated with 100 µg/mL of LY or FITC-label dextran at 37°C for 30 minutes. The black dotted line refers to normal (blank control). (*C*, *D*) The mean fluorescence intensity of LY and FITC-label dextran in MDCK (*C*) and T24 (*D*) cells as determined by flow cytometry with or without additional 10 nM or 5 nM βγ-CAT, respectively. The black dotted line refers to normal (blank control). (*E*-*G*) Localization of βγ-CAT and FITC-label dextran in toad UB epithelial cells (*E*), peritoneal cells (*F*) and MDCK cells (*G*) with or without the treatment with 50 nM (UB cells), 50 nM (peritoneal cells) or 10 nM (MDCK cells) βγ-CAT, respectively, for 15 minutes by immunofluorescence. (Scale bars, 30 µm) The results are reported as mean ± SD of triplicate samples in *A*-*D*. *$P < 0.05$, **$P < 0.01$, ***$P < 0.001$ and ****$P < 0.0001$ by unpaired *t* test. All data are representative of at least two independent experiments.

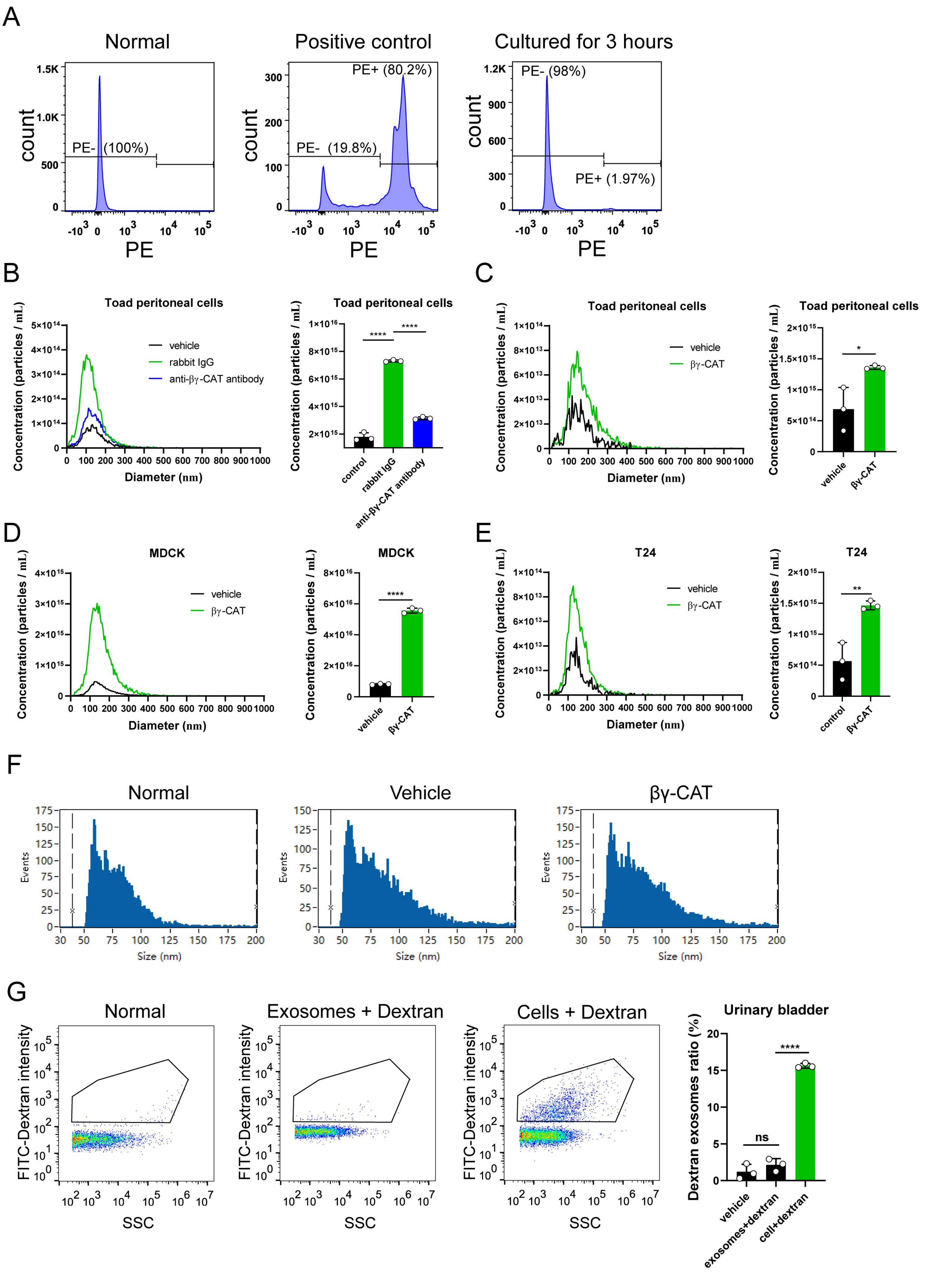


**Fig. S4. βγ-CAT enhances exosome release.** (*A*) The survival rate of toad UB epithelial cells cultured *in vitro* for 3 hours was analyzed by flow cytometry. Digested UB epithelial cells were cultured in isotonic Ringer's solution at room temperature for

3 hours and then the cells were incubated with 500 ng/mL propidium iodide PBS for 10 minutes. (*B*, *C*) NTA analysis of diameter and particle concentration of exosomes isolated from toad peritoneal cells which were cultured in the presence or absence of 50 μg/mL anti-βγ-CAT antibody (*B*) or 50 nM βγ-CAT (*C*) at room temperature for 3 hours. (*D*, *E*) NTA analysis of diameter and particle concentration of exosomes isolated from MDCK (*D*) and T24 (*E*) cells cultured in the presence or absence of 10 nM (MDCK) or 5 nM (T24) βγ-CAT for 3 hours at 37°C, respectively. (*F*) Nanoflow cytometry analysis of the diameter of exosomes derived from toad UB epithelial cells cultured for 3 hours in the presence of 1mg/mL dextran with or without the addition of 50 nM βγ-CAT. (*G*) Comparison of dextran exosome percentage between exosomes cultured directly with 1 mg/mL dextran medium and those collected from UB epithelial cells cultured with 1mg/mL FITC-label dextran medium for 3 hours. The data were obtained by Nanoflow Cytometry and the quantitative result was presented as a bar chart. The results are reported as mean ± SD of triplicate samples in *B*-*E*, *G*. *$P < 0.05$, **$P < 0.01$, ***$P < 0.001$ and ****$P < 0.0001$ by unpaired *t* test. All data are representative of at least two independent experiments.

**Table S1. Sequences of primers used in this study**

| Name | Sequence |
|---|---|
| βγ-CAT-α-Forward | GCTTCCTCTCTGCGTGTGAT |
| βγ-CAT-α-Reverse | GCTTGATAACTGGGTCCCCC |
| βγ-CAT-β-Forward | GCAGCATATGACAGAATTGCATGTCC |
| βγ-CAT-β-Reverse | ACATCCAACTCTTTCTGCAGGGTC |
| β-actin-Forward | GTAGCCCCTGAAGAACACCC |
| β-actin-Reverse | TTGCATGGGGCAGAGCATAA |
